# See and Sequence: Integrating Whole-Genome Sequencing Within the National Antimicrobial Resistance Surveillance Program in the Philippines

**DOI:** 10.1101/808378

**Authors:** Silvia Argimón, Melissa A. L. Masim, June M. Gayeta, Marietta L. Lagrada, Polle K. V. Macaranas, Victoria Cohen, Marilyn T. Limas, Holly O. Espiritu, Janziel C. Palarca, Jeremiah Chilam, Manuel C. Jamoralín, Alfred S. Villamin, Janice B. Borlasa, Agnettah M. Olorosa, Lara F.T. Hernandez, Karis D. Boehme, Benjamin Jeffrey, Khalil Abudahab, Charmian M. Hufano, Sonia B. Sia, John Stelling, Matthew T.G. Holden, David M. Aanensen, Celia C. Carlos, on behalf of the Philippines Antimicrobial Resistance Surveillance Program

**Affiliations:** Centre for Genomic Pathogen Surveillance, Wellcome Genome Campus, Hinxton, UK; Antimicrobial Resistance Surveillance Reference Laboratory, Research Institute for Tropical Medicine, Muntinlupa, Philippines; Brigham and Women’s Hospital, Boston, MA, USA; University of St Andrews School of Medicine, St Andrews, Scotland; Big Data Institute, Li Ka Shing Centre for Health Information and Discovery, University of Oxford, Oxford, UK

## Abstract

Drug-resistant bacterial infections constitute a growing threat to public health globally ^1^. National networks of laboratory-based surveillance of antimicrobial resistance (AMR) monitor the emergence and spread of resistance and are central to the dissemination of these data to AMR stakeholders ^2^. Whole-genome sequencing (WGS) can support these efforts by pinpointing resistance mechanisms and uncovering transmission patterns ^3, 4^. However, genomic surveillance is rare in low- and middle-income countries (LMICs), which are predicted to be the most affected by AMR ^5^. We implemented WGS within the established Antimicrobial Resistance Surveillance Program (ARSP) of the Philippines via ongoing technology transfer, capacity building in and binational collaboration. In parallel, we conducted an initial large-scale retrospective sequencing survey to characterize bacterial populations and dissect resistance phenotypes of key bug-drug combinations, which is the focus of this article. Starting in 2010, the ARSP phenotypic data indicated increasing carbapenem resistance rates for *Pseudomonas aeruginosa*, *Acinetobacter baumannii*, *Klebsiella pneumoniae* and *Escherichia coli*. We first identified that this coincided with a marked expansion of specific resistance phenotypes. By then linking the resistance phenotypes to genomic data, we revealed the diversity of genetic lineages (strains), AMR mechanisms, and AMR vehicles underlying this expansion. We discovered a previously unreported plasmid-driven hospital outbreak of carbapenem-resistant *K. pneumoniae*, uncovered the interplay of carbapenem resistance genes and plasmids in the geographic circulation of epidemic *K. pneumoniae* ST147, and found that carbapenem-resistant *E. coli* ST410 consisted of diverse lineages of global circulation that carried both international and local plasmids, resulting in a combination of carbapenemase genes variants previously unreported for this organism. Thus, the WGS data provided an enhanced understanding of the interplay between strains, genes and vehicles driving the dissemination of carbapenem resistance in the Philippines. In addition, our retrospective survey served both as the genetic background to contextualize local prospective surveillance, and as a comprehensive dataset for training in bioinformatics and genomic epidemiology. Continued prospective sequencing, capacity building and collaboration will strengthen genomic surveillance of AMR in the Philippines and the translation of genomic data into public-health action. We generated a blueprint for the integration of WGS and genomic epidemiology into an established national system of laboratory-based surveillance of AMR through international collaboration that can be adapted and utilized within other locations to tackle the global challenge of AMR.

## Introduction

Antimicrobial resistance (AMR) is an increasingly serious threat to global public health and the economy that requires concerted action across countries, government sectors and non-government organizations ^1^. Without AMR containment, an adverse impact on medical costs, global gross domestic product (GDP), livestock production and international trade is expected by 2050, and the sustainable development goals for 2030 are less likely to be attained ^5^. The Global Action Plan developed by the World Health Organization (WHO) to tackle AMR highlights the need to strengthen our understanding of how resistance develops and spreads, and the underlying resistance mechanisms ^2^. One of the pillars of this objective is the national surveillance system for antimicrobial resistance. An advanced example of a national surveillance system is the Antimicrobial Resistance Surveillance Program (ARSP), the national laboratory-based surveillance of the Philippine Department of Health (DOH). The ARSP began in 1988 with ten tertiary hospitals (sentinel sites) and the Antimicrobial Resistance Surveillance Reference Laboratory (ARSRL) in Metro Manila and has since expanded to 25 sentinel sites and two gonorrhoea surveillance sites in all 17 regions of the country (Supplementary Figure S1). Surveillance encompasses common pathogens of public health importance in the Philippines, where infectious diseases represented nine out of the ten leading causes of morbidity ^6^, and four out of the ten leading causes of infant mortality ^7^ in 2013-2014. Results of culture-based identification and antimicrobial susceptibility are stored centrally at the ARSRL within the clinical microbiology management software WHONET (Figure 1A, ^8^). A report summarizing the resistance trends is published annually and disseminated to local, national and international surveillance stakeholders ^9^. For policy change, surveillance data generated by ARSP is used by the DOH to develop clinical practice guidelines and determine the panel of antibiotics to be included in the national formulary. Furthermore, ARSP has been contributing data to international AMR surveillance programs since the 1980s, including the Global Antimicrobial Surveillance System (GLASS ^10^) since 2015. Importantly, the ARSP data has informed the Philippine National Action Plan to combat AMR, one of the key requirements for alignment with the WHO global action plan.

**Figure 1.**
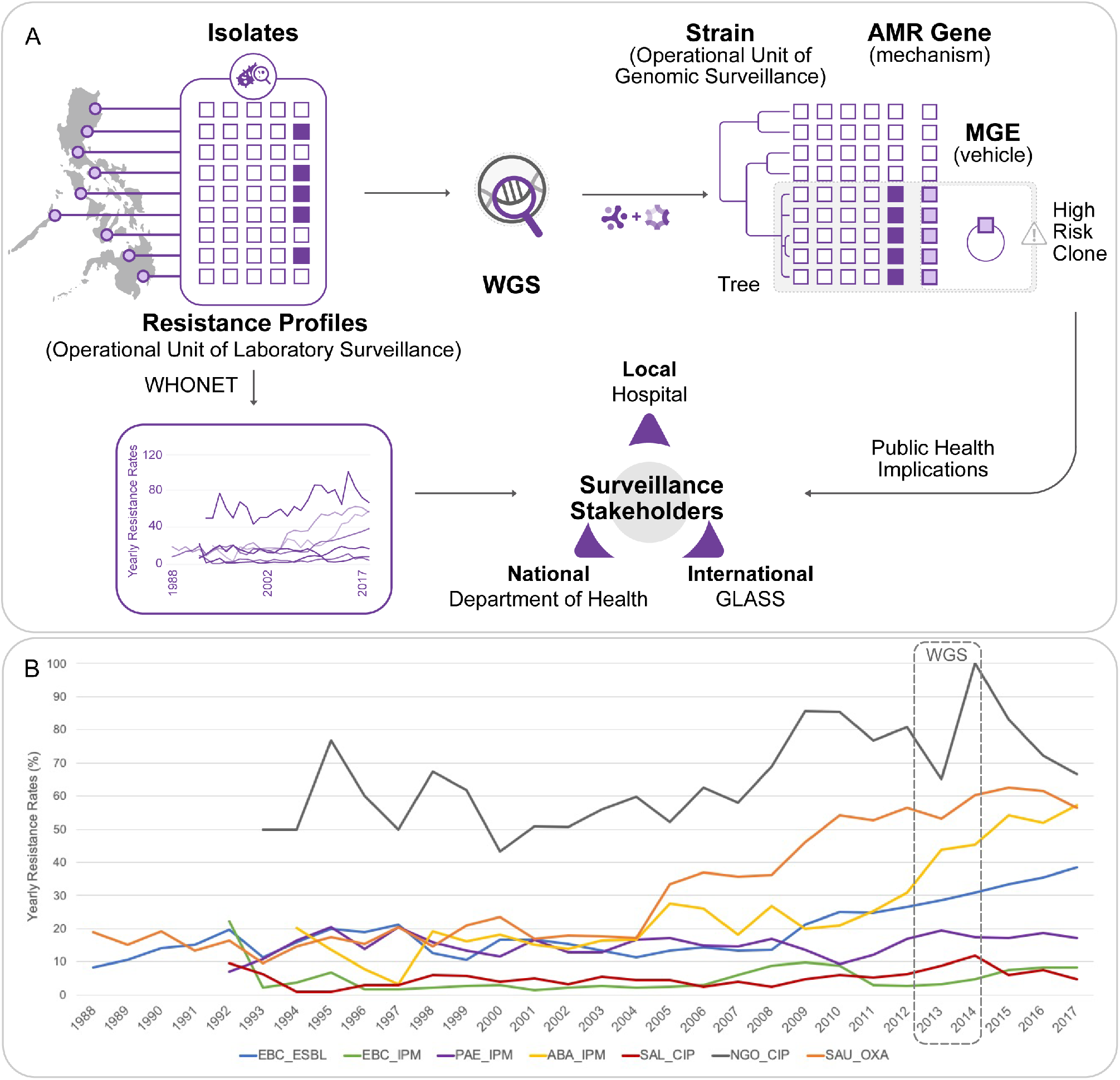
Implementing WGS for AMR surveillance in the Philippines. **A)** ARSP workflow and enhanced detection of high-risk clones by WGS. Isolates collected by sentinel sites are tested for susceptibility to antibiotics (open squares: susceptible, solid squares: resistant). The data are stored as resistance profiles in WHONET and summaries of resistance trends are shared yearly with surveillance stakeholders. Whole-genome sequencing (WGS) of bacterial isolates provides information on genetic relatedness (strains), known AMR mechanisms, and their vehicles for dissemination, allowing us to detect high-risk clones. **B)** Detail of trends in antimicrobial resistance in the Philippines. Yearly resistance rates for key bug-drug combinations based on phenotypic data collected by sentinel sites. EBC: Enterobacteriaceae (*K. pneumoniae*, *E. coli*, *Salmonella enterica*), PAE: *P. aeruginosa*, ABA: *A. baumannii*, but before 2000 *Acinetobacter* spp. SAL: *Salmonella enterica*, SAU: *S. aureus*. ESBL: Extended-spectrum beta-lactamase production suspected, or non-susceptible to the following antibiotics IPM: imipenem, CIP: ciprofloxacin, OXA: oxacillin. Dashed-rectangle labelled with WGS: period covered by the retrospective sequencing survey.

The Philippines has seen a steady increase in resistance rates for several key pathogen-antibiotic combinations in the last ten years, including carbapenem-resistant organisms (Figure 1B). The genetic mechanisms underlying carbapenem resistance, one of the biggest therapeutic challenges in the treatment of antimicrobial resistant infections, include increased upregulation of efflux pumps, decreased uptake by altered expression/loss of porin function, and acquisition of hydrolytic enzymes –carbapenemases ^11^. Whole-genome sequencing (WGS) of bacterial pathogens can identify distinct clonal lineages (strains) on phylogenetic trees, known AMR mechanisms (genes or mutations) and the vehicles for the acquired AMR genes (mobile genetic elements, MGEs) which, in turn, enables enhanced detection and characterization of high-risk clones (Figure 1A). WGS is routinely used for infectious disease epidemiology in several high-income countries around the world, where it has improved outbreak investigations and epidemiological surveillance ^12, 13^ and enhanced our knowledge of the spread of antimicrobial-resistant strains and their resistance mechanisms ^3, 4^. Elucidation of resistance mechanisms and their context can be critical for effective infection control. For example, upregulation of efflux pumps and loss of porin function by mutation are vertically transmitted, while acquired carbapenemases carried in transmissible plasmids or integrative conjugative elements have the potential for horizontal dissemination between strains and species, thus necessitating enhanced infection control measures ^14^. Moreover, diverse carbapenemase classes, variants, flanking MGEs, and associated plasmid incompatibility groups have been described, requiring multiple biochemical and molecular tests for identification. For example, the class B New Delhi metallo-beta-lactamase (NDM) is found worldwide, represented by over ten different variants that are usually upstream of intact or truncated IS*Aba125* elements, within plasmid backbones of different incompatibility groups, across multiple lineages of *E. coli* –as well as of other carbapenem-resistant organisms ^15, 16^. WGS can determine the interplay between these different components, i.e. the gene, the vehicle, and the strain, thus maximizing the epidemiological benefit derived from its cost.

Implementation of WGS within existing national surveillance systems in LMICs has the potential to enhance local prevention and control of resistant infections in a sustainable and equitable manner. Integration of WGS into routine surveillance of AMR can be facilitated by international collaboration focused on transfer of expertise and ownership. Our collaboration (See and Sequence, Figure 1A) aimed to implement WGS within the ARSP via a multi-faceted approach that included a large initial retrospective sequencing survey, technology transfer, utilization of user-friendly web applications, and capacity building in laboratory and bioinformatic procedures for local prospective sequencing. We linked what has traditionally been used as the operational unit of laboratory surveillance, the resistance profile, with the operational unit of genomic epidemiology now provided by WGS, genetic relatedness. Here we provide exemplars from the retrospective sequencing survey that highlight how the granular view of strain-gene-vehicle in carbapenem-resistant populations at the local, national and global operational scales can be leveraged for surveillance of AMR and public health.

## Results

The Philippine Antimicrobial Resistance Surveillance Program (ARSP) collects bacterial isolates and stores the associated clinical and epidemiological data in the WHONET software ^8^. Antibiograms are stored as resistance profiles, which represent the diversity of AMR phenotypes in the country. In parallel to local WGS capacity building, we conducted a large retrospective sequencing survey of eight bacterial pathogens to provide genomic context for local prospective surveillance. Here, we focus on the carbapenem-resistant organisms, which have been classified as of critical priority for the development of new antibiotics by the World Health Organization (WHO).

### Operational unit of laboratory surveillance - Routine laboratory surveillance highlights the increasing burden of specific resistance phenotypes

Carbapenem-resistant *P. aeruginosa, A. baumannii, K. pneumoniae* and *E. coli* were all first isolated by ARSP between 1992 and 1994. Since the year 2000 resistance rates to imipenem and meropenem have remained below 30% for all organisms except for *A. baumannii*, which has seen a steady rise in resistance rates between 2009 and 2017, reaching 56% (Figure 2A). This coincides with the expansion of two resistance profiles (RP), first a striking expansion of true-XDR and possible-PDR RP-1, and later of true-XDR RP-2 (full profiles can be found in Figure 2B). Yearly resistance rates for *P. aeruginosa* have oscillated between 10-25% since the year 2000, and have doubled between 2010 and 2013 (Figure 2A). Two resistance profiles have seen the largest combined expansion between 2011 and 2013, possible-PDR RP-3, and true-XDR RP-4 (Figure 2B). The yearly resistance rates to imipenem for Enterobacteriaceae were approximately 3% in 2011 but have since steadily increased to 5% for *E. coli* and 11% for *K. pneumoniae*, respectively (Figure 2A). This coincides with the expansion of three possible-XDR resistance profiles, RP-5, RP-6, and RP-7 (Figure 2B). The observed expansion of specific resistance profiles could be driven by the rapid dissemination of a discrete genetic lineage (strain) carrying AMR determinants or, alternatively, of transmissible MGEs, which shuttle resistance determinants across different genetic lineages.

**Figure 2.**
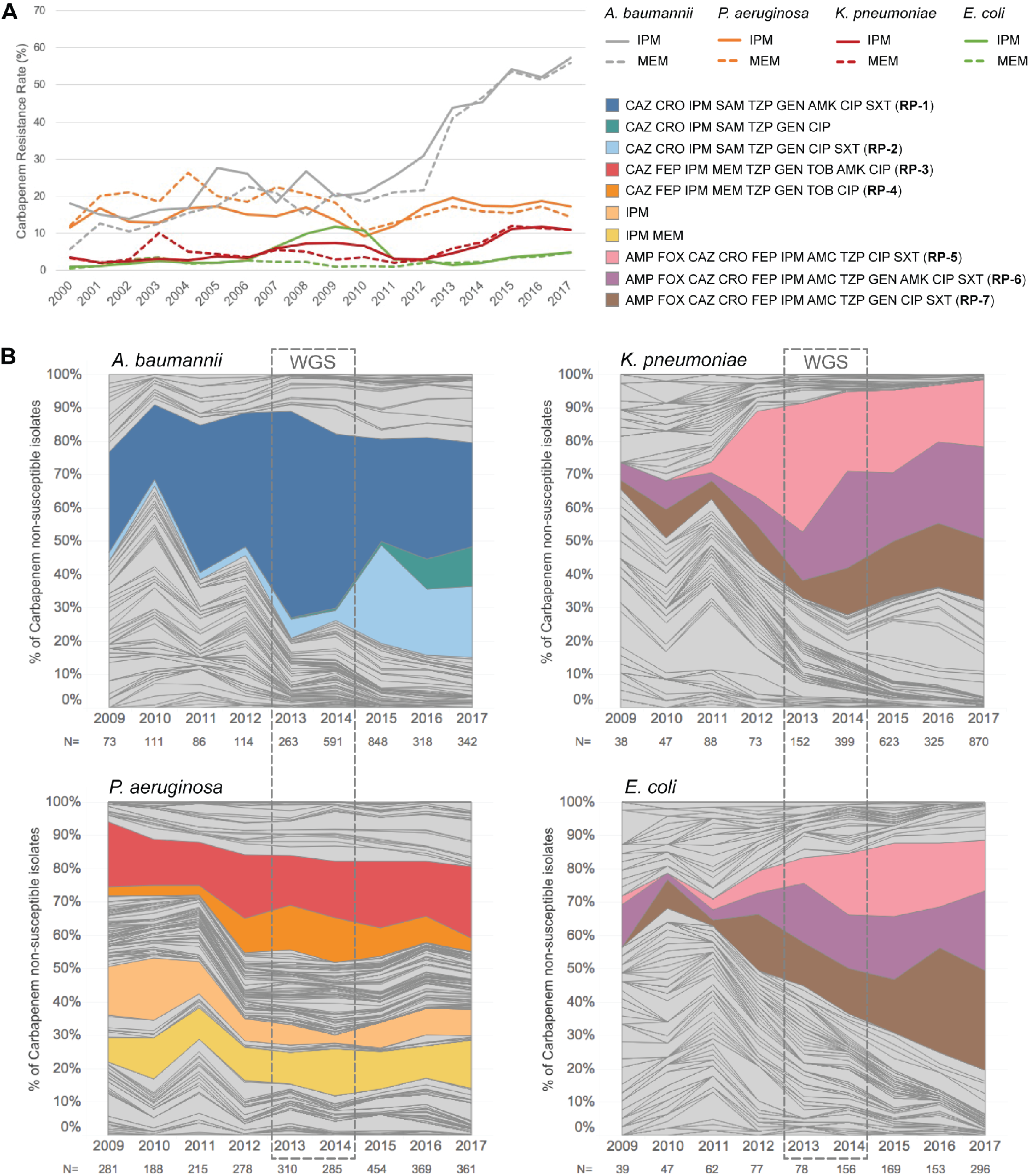
Temporal dynamics of the resistance profiles from carbapenem non-susceptible isolates. **A)** Yearly carbapenem resistance rates (IPM: imipenem, MEM: meropenem) for *P. aeruginosa*, *A. baumannii*, *E. coli* and *K. pneumoniae*. **B)** Relative abundance of resistance profiles with resistance to carbapenems (imipenem and/or meropenem). The three-letter code of an antibiotic indicates that the isolate is non-susceptible (resistant or intermediate) to the antibiotic. Only carbapenem non-susceptible isolates with complete susceptibility data were included, as indicated by the numbers under the the x-axis (N). Dashed-rectangle labelled with WGS: period covered by the retrospective sequencing survey.

### Operational unit of genomic surveillance - WGS reveals the genetic diversity and AMR mechanisms underpinning carbapenem resistance phenotypes

The sequences of 805 genomes linked to WHONET data were obtained from isolates belonging to the four bacterial pathogens in the WHO critical list (Table 1), which were collected in 2013 and 2014 by between thirteen to eighteen ARSP sites. The isolates sequenced were non-susceptible to carbapenems and/or to extended-spectrum cephalosporins (Table 1), with the exception of *P. aeruginosa*, for which susceptible isolates were available and also sequenced.

**Table 1.** Organisms from the retrospective sequencing survey described in this study. The number of draft genomes obtained, and the number of sentinel sites they represent are indicated. Key resistances were used to prioritize the isolates for WGS as described in the methods. Antibiotics tested by ARSRL for each organism are abbreviated according to their WHONET codes: AMC: amoxicillin/clavulanic acid, AMK: amikacin, AMP: ampicillin, CAZ: ceftazidime, CIP: ciprofloxacin, CRO: ceftriaxone, GEN: gentamicin, FEP: cefepime, FOX: cefoxitin, IPM: imipenem, MEM: meropenem, SAM: ampicillin/sulbactam, SXT: trimethoprim/sulfamethoxazole, TOB: tobramycin, TZP: piperacillin/tazobactam.

| Organism (code) | Key Resistance(s) | Antibiotics tested by ARSP | Genomes | Sentinel Sites |
| --- | --- | --- | --- | --- |
| *K. pneumoniae* (Kpn) | Carbapenems, ESBL-producing | AMP, FOX, CAZ, CRO, FEP, IPM, AMC, TZP, GEN, AMK, CIP, SXT | 340 | 18 |
| *E. coli* (Eco) | Carbapenems, ESBL-producing | AMP, FOX, CAZ, CRO, FEP, IPM, AMC, TZP, GEN, AMK, CIP, SXT | 181 | 17 |
| *P. aeruginosa* (Pae) | Carbapenems | CAZ, FEP, IPM, MEM, TZP, GEN, TOB, AMK, CIP | 176 | 17 |
| *A. baumannii* (Aba) | Carbapenems | CAZ, CRO, IPM, SAM, TZP, GEN, AMK, CIP, SXT | 108 | 13 |

The distribution of the carbapenem resistance profiles highlighted in Figure 2 was not concordant with the major clades observed on the phylogenetic trees (Figure 3). Instead, the same resistance profile was usually observed in multiple genetic lineages across the tree. Yet, the distribution of the multi-locus sequence types (STs), a molecular genotyping method inferred from the sequences of seven house-keeping genes, was largely concordant with the major clades observed on the trees inferred from whole-genome single-nucleotide polymorphisms (SNPs) (Figure 3), confirming that the homoplasic distribution of resistance profiles was not an artefact of the tree topology. The distribution of the same carbapenem resistance profiles across multiple genetic lineages was observed both at the national and local levels (Supplementary Figure S2), as the number of ST assignments increased with the number of isolates representing a resistance profile. Thus, the genomic data suggests that expansion of specific carbapenem resistance profiles in the Philippines is driven by the horizontal dissemination of resistance determinants across diverse genetic backgrounds, rather than by the sole expansion of single resistant genetic clones, and that resistance to multiple antibiotics is acquired *en bloc* via MGEs, as previously observed for carbapenem-resistant organisms ^16^.

**Figure 3.**
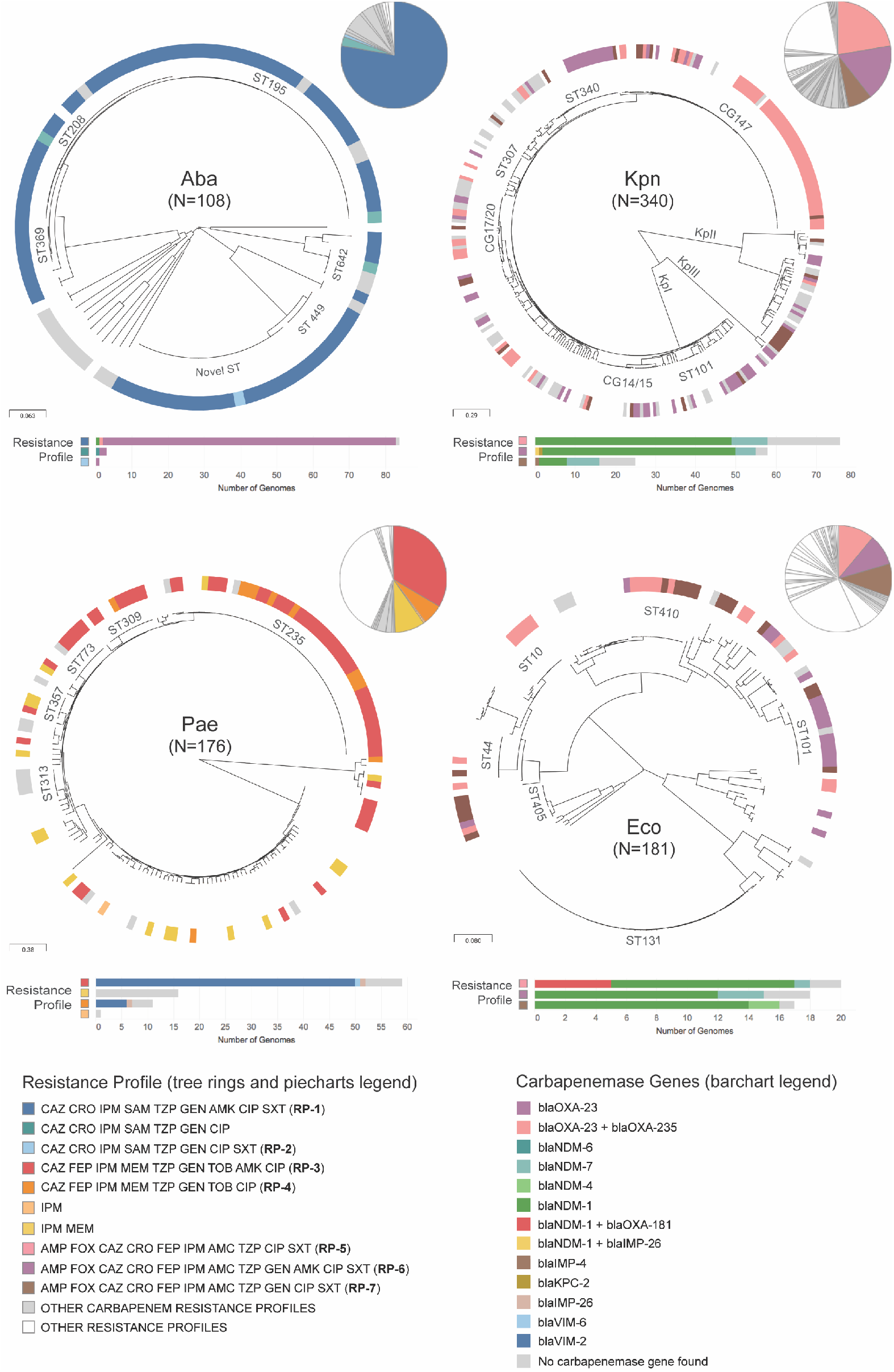
Resistance profiles are not associated with genetic lineages in carbapenem non-susceptible organisms. Phylogenetic trees showing the major lineages of *A. baumannii* (Aba), *P. aeruginosa* (Pae), *K. pneumoniae* species complex (Kpn), and *E. coli* (Eco), indicating the position of select STs and clonal groups (CGs). KpI: *K. pneumoniae sensu stricto*, KpII: *K. quasipneumoniae*, KpIII: *K. variicola*. Tree ring: Select carbapenem resistance profiles from Figure 2 are shown in colour. The remaining carbapenem resistance profiles are shown in light grey. Other resistance profiles (susceptible to carbapenems) are shown in white for simplicity. The pie charts show the relative abundance of the resistance profiles in the retrospective collection of sequenced genomes. The bar charts show the distribution of carbapenemase genes across the key resistance profiles. The data are available at https://microreact.org/project/ARSP_ABA_2013-14 (Aba), https://microreact.org/project/ARSP_PAE_2013-14 (Pae), https://microreact.org/project/ARSP_KPN_2013-14 (Kpn), and https://microreact.org/project/ARSP_ECO_2013-14 (Eco).

Carbapenemases are a diverse collection of hydrolytic enzymes ^16^, and we identified representatives of class A, B and C carbapenemases in the Philippines (Supplementary Figure S3). Class B NDM-1 and Verona integron-borne metallo-beta-lactamase VIM-2 were the most prevalent and geographically disseminated in the Enterobacteriaceae and *P. aeruginosa,* respectively, while the same was true for class D OXA-23 in *A. baumannii*. Importantly, different carbapenemase genes/variants (or combinations thereof) were found underlying the same resistance profiles in all four organisms (Figure 3).

### Scale of surveillance – Local: WGS reveals a plasmid driven hospital outbreak of carbapenem-resistant K. pneumoniae

Fifty-seven percent (N=33) of the *K. pneumoniae* genomes with the possible-XDR resistance phenotype “AMP FOX CAZ CRO FEP IPM AMC TZP GEN AMK CIP SXT” (RP-6) were isolated from a single hospital (MMH) but belonged to 12 different STs, with almost half of them (N=15) placed within ST340 (Supplementary Figure S4A). Phylogenetic analysis of these 15 genomes in the wider context of ST340 indicated three main lineages, with the 15 possible-XDR isolates from MMH forming a tight cluster (clade III, Figure 4A). The shorter branch lengths in the imipenem-resistant clade III (6.5±3.7 pairwise SNP differences) compared to the imipenem-susceptible clade II (26.8 ±6.9 pairwise SNP differences, Mann-Whitney U test z-score −6.265, p-value= 3.71×10^−10^, Supplementary Figure S4B), and the isolation dates spanning twelve months, suggest a rapid expansion of this clone, which coincides with the acquisition of the *bla*_NDM-1_ gene (Figure 4A).

**Figure 4.**
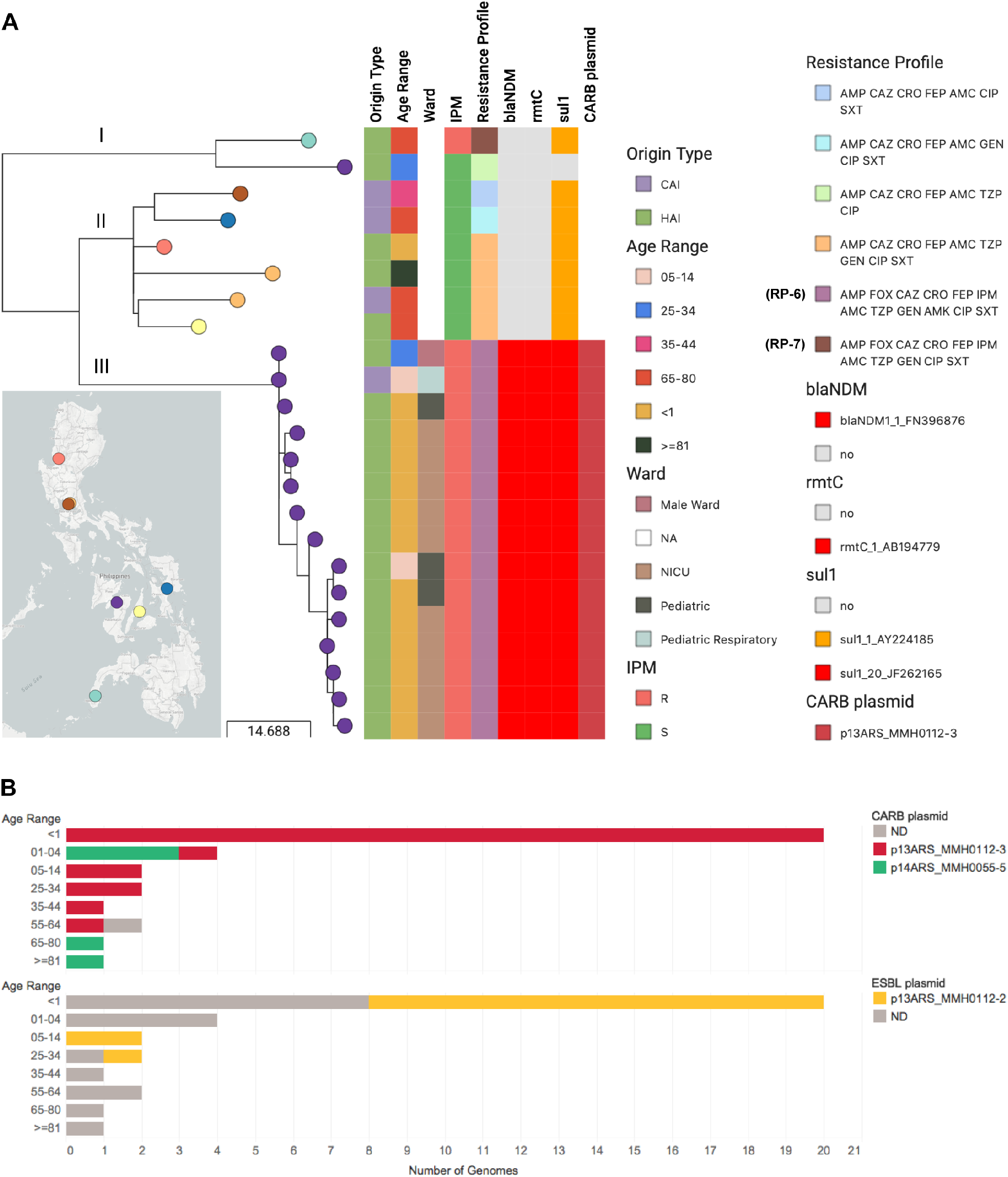
WGS reveals a previously undetected, plasmid-driven outbreak of *K. pneumoniae* ST340. **A)** Phylogenetic tree and linked epidemiological and genotypic data of 24 retrospective ST340 genomes. This interactive view is available at https://microreact.org/project/ARSP_KPN_ST340_2013-14/ac2a0920. Maximum-likelihood tree inferred from 196 SNP positions identified by mapping the genomes to reference CAV1217 (GCA_001908715), and masking regions corresponding to mobile genetic elements and recombination. The full data are available at https://microreact.org/project/ARSP_KPN_ST340_2013-14. **B)** Distribution of 33 retrospective isolates from hospital MMH with resistance profile RP-6 by patient age group, with the distribution of STs (top panel) and plasmids with carbapenemases genes (bottom panel) indicated by the different colours. Short reads of the 33 isolates were mapped to the p13ARS_MMH0112-3 and the p14ARS_MMH0055-5 sequences and a plasmid match was counted when the reads covered at least 95% of the sequence length with at least 5x depth of coverage.

Epidemiological data showed that the samples were mostly hospital-acquired (N=14), and from neonates (N=12) (Figure 4A). These observations triggered a retrospective epidemiological investigation that revealed that 10 of the isolates originated from patients of the neonatal intensive care unit (NICU), all of which had umbilical catheters and were on mechanical ventilators. The remaining cases were either from paediatric wards (N=4) or the male ward (N=1), suggesting wider dissemination of this high-risk clone within the hospital environment.

The genetic diversity underlying the 33 possible-XDR isolates from MMH (12 STs) prompted us to investigate the hypothesis of dissemination of carbapenem resistance in the hospital by a plasmid carrying *bla*_NDM-1_. We identified a 101,540 bp IncFII plasmid with *bla*_NDM-1_, *rmtC*, and *sul1* (p13ARS_MMH0112-3, Table 2 and Supplementary Results and Figure S5) in a representative isolate from ST340 clade III. In addition, we found the entire p13ARS_MMH0112-3 plasmid sequence (i.e. ≥95% of the length) represented in 27 genomes from 9 different sequence types, including 20 samples from 6 different sequence types isolated from patients under 1 year old (Figure 4B).

**Table 2.**
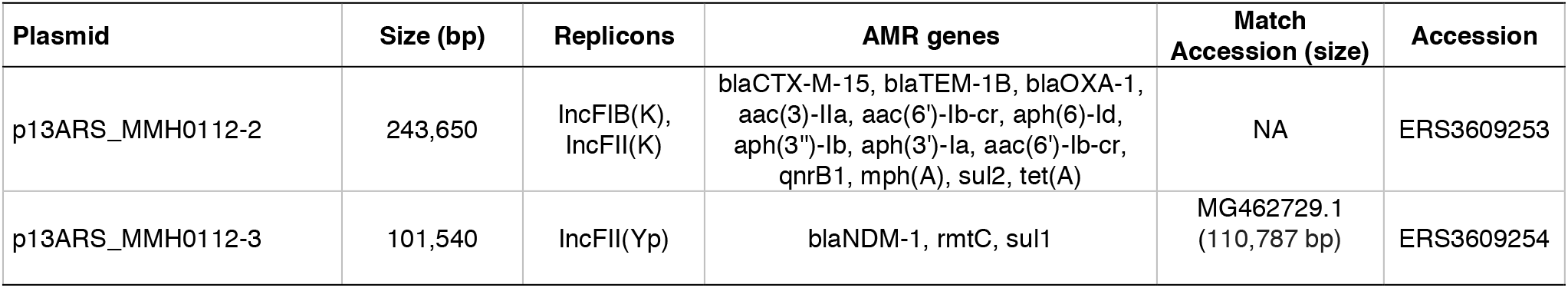
Plasmids carrying carbapenemase genes and other resistance genes in *K. pneumoniae* ST340 strain 13ARS_MMH0112 from the Philippines. The replicons and AMR genes were identified with the plasmidFinder and Resfinder databases, respectively. Match Accession indicates the accession number when at least one match of more than 90% query sequence coverage and 99% identity was found in the NCBI nucleotide database.

| Plasmid | Size (bp) | Replicons | AMR genes | Match Accession (size) | Accession |
| --- | --- | --- | --- | --- | --- |
| p13ARS_MMH0112-2 | 243,650 | IncFIB(K), IncFII(K) | blaCTX-M-15, blaTEM-1B, blaOXA-1, aac(3)-IIa, aac(6)-Ib-cr, aph(6)-Id, aph(3'')-Ib, aph(3')-Ia, aac(6)-Ib-cr, qnrB1, mph(A), sul2, tet(A) | NA | ERS3609253 |
| p13ARS_MMH0112-3 | 101,540 | IncFII(Yp) | blaNDM-1, rmtC, sul1 | MG462729.1 (110,787 bp) | ERS3609254 |

Altogether, our findings suggest that the burden of carbapenem-resistant *K. pneumoniae* infections in hospital MMH was largely linked to plasmid p13ARS_MMH0112-3 with *bla*_NDM-1_, circulating within diverse genetic lineages, and leading to outbreaks in high-risk patient populations. Hospital authorities were informed of these findings, and measures for infection control were implemented, including designating a separate multi-drug resistance organism (MDRO) room for cohorting, active surveillance upon identification of any new carbapenem-resistant *K. pneumoniae* from the NICU, and referral of any new carbapenem-resistant *K. pneumoniae* from the NICU to ARSRL for sequencing.

### Scale of surveillance - National: WGS reveals the interplay of carbapenem resistance genes and plasmids in the regional circulation of a successful K. pneumoniae lineage

The possible-XDR resistance phenotype “AMP FOX CAZ CRO FEP IPM AMC TZP CIP SXT” (RP-5) was represented by 76 *K. pneumoniae* isolates from 14 different STs. Seventy-one percent (N=54) belonged to ST147, an international epidemic clone that was found in 11 sentinel sites in all 3 island groups in the Philippines. The phylogeny of the ST147 genomes showed that carbapenem non-susceptible isolates (78·8%, N=63) were found in clades arising from 3 out of 4 deep branches of the tree (Figure 5A), which represent separate groups of the global population (Supplementary Figure S6A). Carbapenem resistance coincided with the presence of *bla*_NDM-1_ in clade IV genomes, while clade III-B shows geographically distinct clusters with either *bla*_NDM-1_ or *bla*_NDM-7_ (Figure 5A and Supplementary Results and Figure S7).

**Figure 5.**
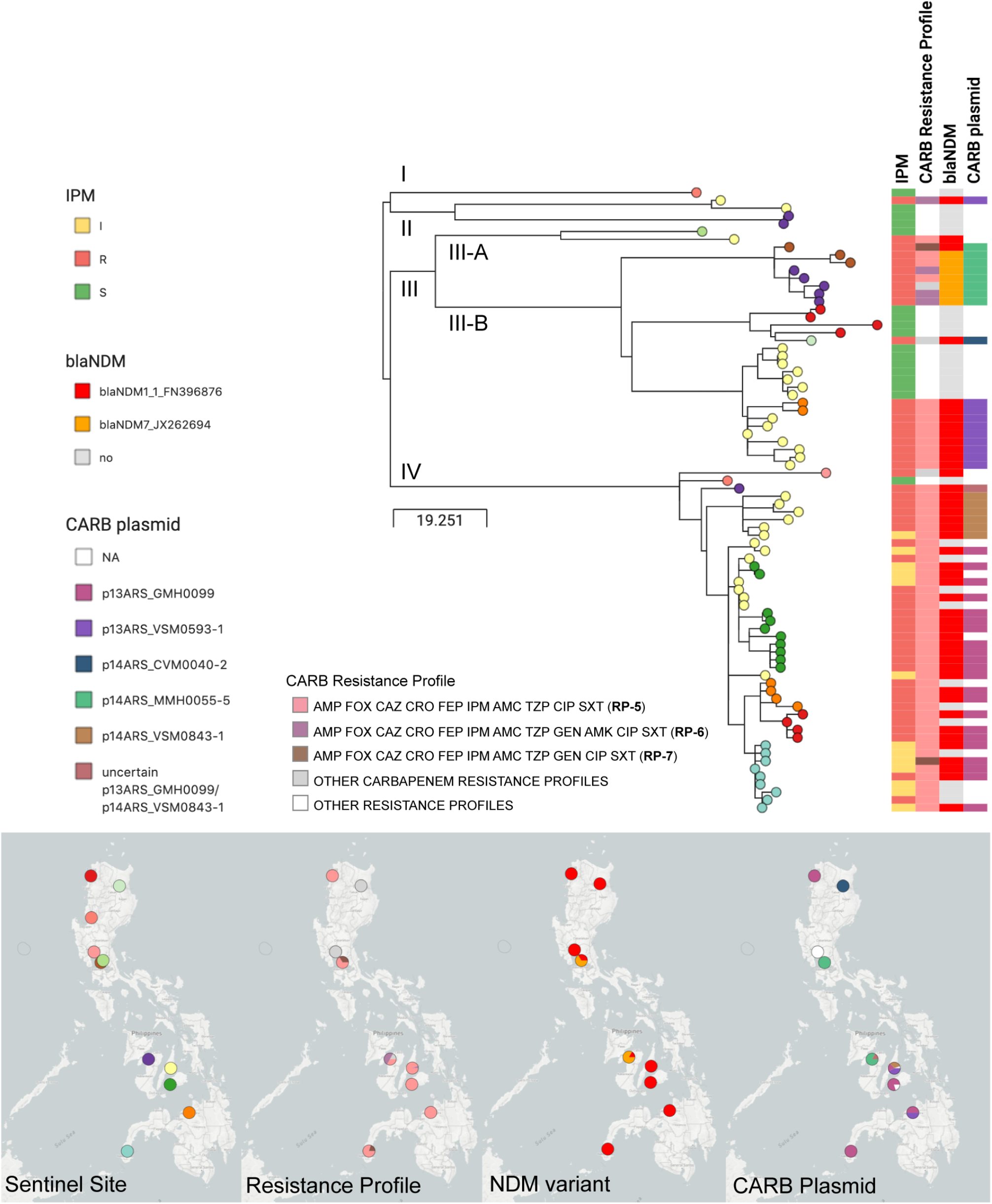
WGS reveals the circulation patterns of *K. pneumoniae* ST147 in the Philippines. Phylogenetic tree and linked epidemiological and genotypic data of 80 retrospective ST147 genomes. This interactive view is available at https://microreact.org/project/ARSP_KPN_ST147_2013-14/4c33ace7. Maximum-likelihood tree inferred from 809 SNP positions identified by mapping the genomes to reference MS6671 (LN824133.1) and masking regions corresponding to mobile genetic elements and recombination. The distribution of plasmids with carbapenemases genes was inferred by mapping the short reads of the genomes to the complete plasmid sequences, and a match was counted when the reads covered at least 95% of the sequence length with at least 5x depth of coverage. The full data are available at https://microreact.org/project/ARSP_KPN_ST147_2013-14.

Plasmid sequences obtained from isolates representing different carbapenem-resistant clusters in clades III and IV showed that the NDM genes were carried within different variants of the insertion sequence IS*Aba*125 on different plasmid backbones, two of which showed high sequence similarity to international plasmids (Table 3 and Supplementary Figures S5B, S6B and S6C). The distribution of plasmids harbouring *bla*_NDM-1_ and *bla*_NDM-7_ matched the strong phylogeographic signal in the terminal branches of the tree (Figure 5A). Within clade III-B, a cluster of genomes from sentinel sites MMH and STU were characterized by plasmid p14ARS_MMH0055-5 carrying *bla*_NDM-7_, while another cluster of genomes from VSM and NMC were distinguished by plasmid p13ARS_VSM0593-1 with *bla*_NDM-1_. Plasmids p13ARS_GMH0099 and p14ARS_VSM0843-1, both carrying *bla*_NDM-1_ were found in different subclades of clade IV, one of local circulation in hospital VSM, and another one showing regional expansion across 5 sentinel sites (Figure 5A). Taken together, the phylogeographic signal (strains) in combination with the distribution of *bla*_NDM_ variants (mechanism) and plasmids (vehicle), revealed local and regional patterns of circulation of carbapenem non-susceptible ST147 within the Philippines, which could not be identified based solely on the resistance profiles or MLST information. Furthermore, comparison with global ST147 genomes indicated that clade IV may be unique to the Philippines (Supplementary Results and Figure S6A).

**Table 3.** Plasmids carrying NDM genes in *K. pneumoniae* ST147 representative isolates from the Philippines. The replicons and AMR genes were identified with the plasmidFinder and Resfinder databases, respectively. Match Accession indicates the accession number when at least one match of more than 90% query sequence coverage and 99% identity was found in the NCBI nucleotide database.

| Plasmid | Size (bp) | Replicons | AMR genes | Match Accession (size) | Accession |
| --- | --- | --- | --- | --- | --- |
| p14ARS_CVM0040-2 | 110,787 | IncFII(Yp) | blaNDM-1, rmtC, sul1 | MG462729.1 (110,787 bp) | ERS3609255 |
| p13ARS_GMH0099 | 273,158 | IncA/C2, IncR, IncFIA(HI1) | blaNDM-1, blaCTX-M-15, blaOXA-1, blaOXA-10, blaTEM-1B, aph(3')-VI, aph(3')-Ia, aadA1(x2), aac(6)-Ib-cr, aph(6)-Id, strA, strB, qnrS1, mph(A), erm(42), ere(A), floR, cmlA1(x2), ARR(x2), sul1(x3), sul2, dfrA14 | NA | ERS3609256 |
| p13ARS_VSM0593-1 | 181,414 | IncA/C2 | blaNDM-1, aph(3')-Ia, mph(A), cmlA1, floR, sul1, sul2, qnrS1, ARR-3, | NA | ERS3609257 |
| p14ARS_MMH0055-5 | 44,885 | IncX3 | blaNDM-7 | KP776609.1 (45,122 bp) | ERS3609258 |
| p14ARS_VSM0843-1 | 95,284 | IncR, IncFIA(HI1) | blaNDM-1, blaCTX-M-15, blaTEM-1B, aph(3')-VI, aph(6)-Id, aph(3')-Ib, aph(3')-VI, aadA1, cmlA1, sul1, sul2, ARR-3, | NA | ERS3609259 |

**Table 4.** Plasmid repertoire of carbapenem-resistant *E. coli* strain 14ARS_NMC0074. The replicons and AMR genes were identified with the plasmidFinder and Resfinder databases, respectively. Match Accession indicates the accession number when at least one match of more than 90% query sequence coverage and 99% identity was found in the NCBI nucleotide database.

| Plasmid | Size (bp) | Replicons | AMR genes | Match Accession (size) | Accession |
| --- | --- | --- | --- | --- | --- |
| p14ARS_NMC0074-2 | 146,817 | IncA/C2 | blaNDM-1, blaOXA-10, aadA1, aph(3')-VI, aph(3')-Ia, mph(A), cmlA1, floR, sul1, sul2, dfrA14, qnrS1, ARR-2 | NA | ERS3609260 |
| p14ARS_NMC0074-5 | 51,479 | IncX3 | blaOXA-181, qnrS1 | CP024806.1 (51,479 bp) | ERS3609263 |
| p14ARS_NMC0074-4 | 81,712 | IncFII/IncFIA | blaCTX-M-15, blaTEM-1B, blaOXA-1, aadA5, aac(6')-Ib-cr, sul1, tet(B), dfrA17 | CP024805.1 (111,310 bp) | ERS3609262 |
| p14ARS_NMC0074-3 | 82,418 | IncFII | blaTEM-1, mph(A), erm(B) | NA | ERS3609261 |
| p14ARS_NMC0074-6 | 2,088 | Col(BS512) | - | CP024803.1 (2,088 bp) | ERS3609264 |

### Scale of surveillance - International: First report of a high-risk clone of E. coli ST410 carrying bla_NDM-1_ and bla_OXA-181_

Recent reports of *E. coli* ST410 international high-risk clones carrying class D and class B carbapenemases ^17, 18^ are particularly alarming in light of their wide geographic distribution and the broad variety of niches this clone can occupy ^19, 20^. ST410 was the second most prevalent ST in the retrospective collection of *E. coli* (13·2%, N=24), encompassing a large proportion of imipenem-resistant isolates (N=15), and the three expanding possible-XDR resistance profiles RP-5,6,7 (Supplementary Figure S8). The phylogenetic tree of 24 ST410 genomes showed that isolates with the possible-XDR profiles clustered within one clade (Figure 6A), which can be further delineated by the distribution of carbapenemase genes and variants into three different clones carrying *bla*_NDM-4_ (N=1), *bla*_NDM-1_ (N=8), or both *bla*_NDM-1_ and *bla*_OXA-181_ (N=5). The phylogenetic analysis with global *E. coli* ST410 genomes confirmed that these are independent clones, as they were found interspersed with international genomes within a major high-risk lineage (Figure 6B and Supplementary Results).

**Figure 6.**
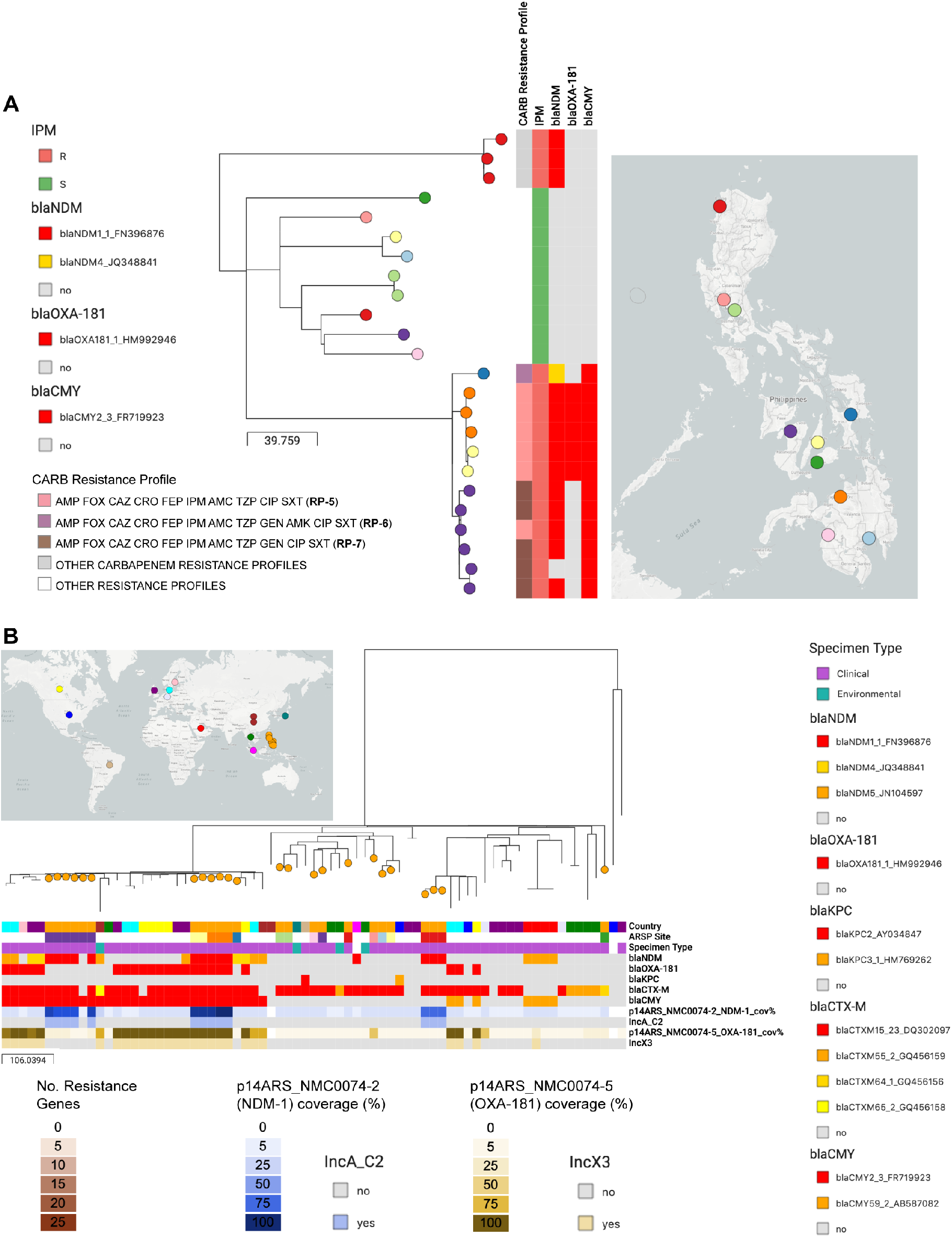
Phylogeographic analysis of *E. coli* ST410 from the Philippines. **A)** Phylogenetic tree and linked epidemiological and genotypic data of 24 retrospective ST410 genomes. This interactive view is available at https://microreact.org/project/ARSP_ECO_ST410/088ba65b. **B)** Philippine isolates (orange nodes) in global context. This interactive view is available at https://microreact.org/project/ARSP_ECO_ST410_GLOBAL/a2286fa8. The maximum-likelihood trees were inferred from 703 (A) and 2851 (B) SNP positions, respectively, identified by mapping the genomes to reference AMA1167, and masking regions corresponding to mobile genetic elements and recombination. The distribution of plasmids with carbapenemases genes in (B) was inferred by mapping the short reads of the genomes to the complete plasmid sequences, and a match was counted when the reads covered at least 95% of the sequence length with at least 5x depth of coverage. The full data are available at https://microreact.org/project/ARSP_ECO_ST410 (A) and https://microreact.org/project/ARSP_ECO_ST410_GLOBAL (B).

*K. pneumoniae* carrying *bla*_NDM-1_ and *bla*_OXA-181_ have been previously reported ^21, 22^, as well as *E. coli* ST410 harbouring *bla*_NDM-5_ and *bla*_OXA-181_ from Egypt ^23^, Denmark, and the UK ^18^, but, to our knowledge, this is the first report of *E. coli* carrying both *bla*_NDM-1_ and *bla*_OXA-181_, which is likely to have disseminated between two sentinel sites (NMC and VSM). We identified five plasmids in the representative strain 14ARS_NMC0074 (Table 3, and Supplemental Figure S9). The IncX3 plasmid carrying *bla*_OXA-181_ (p14ARS_NMC0074-5) was identical to plasmid pAMA1167-OXA-181 isolated previously from an *E. coli* ST410 strain ^24^ (Supplemental Figure S9). Mapping short reads to p14ARS_NMC0074-5 showed that this plasmid is the main vehicle of *bla*_OXA-181_ in the Philippines, as in the international genomes (Figure 6B, ^18^). However, we did not identify any plasmids similar to the IncA/C2 plasmid with *bla*_NDM-1_ (p14ARS_NMC0074-2), though it shared approximately 90% of its backbone with the IncA/C2 plasmid described above from *K. pneumoniae* ST147 strain 13ARS_VSM0593 (Supplemental Figure S9). This plasmid backbone was found in *E. coli* ST410 genomes from the Philippines, but not in international genomes (Figure 6B).

Altogether, our results show that that the Philippine *E. coli* ST410 genomes represent diverse lineages of the global circulating population, with evidence of frequent international introductions. These lineages are characterized by a diverse repertoire of carbapenemase genes and variants, amassed via a combination of conserved international plasmids and locally-circulating plasmids.

## Discussion

National networks of laboratory-based surveillance are a key pillar within the Global Action Plan to combat AMR, with the reference laboratory playing a central role in the dissemination of surveillance data to local, national and international stakeholders. The Philippines ARSP surveillance data has shown increasing resistance trends for key bug-drug combinations (Figures 1B and 2A). In this study, the analysis of the dynamics of resistance profiles showed the concomitant expansion of specific resistance phenotypes (Figure 2B). By complementing laboratory data with WGS and linking the operational units of laboratory and genomic surveillance, we revealed a diversity of genetic lineages, AMR mechanisms, and vehicles underlying the expansion of carbapenem resistance phenotypes (Figures 1A and 3). The combination of these three components vastly improved our understanding of the routes of dissemination of carbapenem resistance in the Philippines at different geographical scales, from an plasmid-driven local outbreak of *K. pneumoniae* ST340 (Figure 4), to the expansion of a *K. pneumoniae* ST147 clone carrying NDM-1 in an IncA/C2 plasmid without any close match in international genomes (Figure 5), to the independent introductions of international epidemic clone *E. coli* ST410 that can acquire plasmids of local circulation (Figure 6).

A crucial aspect of AMR surveillance is the detection of the emergence and spread of high-risk clones. In traditional laboratory surveillance, resistance profiles can be analysed with spatiotemporal algorithms to detect hospital outbreaks ^25^ or dissemination of phenotypic subpopulations between hospitals ^26^. Cluster detection based on resistance profiles depends on consistent and complete antibiograms within and across sentinel sites. During the course of our project it became apparent that there were inconsistencies in the panels of antibiotics tested by the sentinel sites (missing data or different antibiotics tested), which led to the exclusion of samples from the analysis of the dynamics of resistance profiles (Figure 2). These inconsistencies would also hamper cluster detection based on resistance profiles, or the early detection of novel emerging resistance profiles that could act as indicators of novel high-risk clones. The joint discussions of these observations by the two collaborating teams led to an effort to reinforce the standardization of antibiotic testing across the ARSP sentinel sites, which was coordinated by the ARSRL.

Even with complete and comprehensive susceptibility testing, resistance profiles can only provide a coarse view of the spread of AMR, and sometimes lead to clusters of disparate genetic relatedness (Figure 3 and Supplementary Figure 2). Integration of WGS into laboratory-based AMR surveillance can substantially improve the detection of high-risk clones by providing a high-resolution picture of genetic lineages (strains), AMR mechanisms (genes/mutations) and vehicles (MGEs, Figure 1A). We identified high-risk clones within known international epidemic clones at the local, national, and international scales by linking clonal relatedness, geographic clustering, epidemiological data, and gene and plasmid content with interactive web tools ^27^.

At the local scale, we identified a plasmid-driven NICU outbreak of carbapenem-resistant *K. pneumoniae*. The epidemiological data captured in WHONET was key to support the interpretation of the genomic findings, triggering a retrospective investigation. This previously undetected outbreak was traced to an IncFII plasmid carrying *bla*_NDM-1_ in the genetic context of ST340 (Figure 4A). Yet, the endemic IncFII plasmid was found across multiple wards, and genetic backgrounds (STs, Figure 4B), indicating a major role in the burden of carbapenem resistance *K. pneumoniae* in this hospital. A second IncFIB-IncFII plasmid found only within the ST340 lineage might have contributed to the persistence and transmission of this clone, in particular within the NICU, by carrying several genomic islands with a role in survival in the host and in the environment (Supplementary Figure S5). ST340 is a member of the drug-resistant clonal complex 258 and has been reported to cause outbreaks worldwide ^28^. Our findings were disseminated to the local stakeholders (i.e., hospital) via a forum with NICU staff, paediatricians and ARSRL representatives. Infection control strategies were implemented, which ultimately bolstered the infection control team of this hospital. Control of healthcare-associated infections is crucial to containing the spread of antimicrobial resistance ^29^. The roadmap from sequence data to actionable data for infection control and prevention within a hospital has been mapped by multiple use cases ^30, 31^ and supported by recent studies on implementation and cost-effectiveness in high-resource settings ^32, 33^. This study makes a case for extending this roadmap to LMICs that have infection control capacity in place.

At the national scale, the combined information on strain-gene/variant-vehicle shows a granular picture of carbapenem-resistant *K. pneumoniae* ST147 that uncovers the geographic distribution of high-risk clones. ST147 is an international epidemic clone that causes carbapenem-resistant infections mediated by NDM, OXA-48-like and VIM carbapenemases ^28, 34^. However, ST147 was not reported in a recent study of *K. pneumoniae* from seven healthcare facilities across South and Southeast Asia ^35^, which did not include the Philippines. Within the seemingly well-established genetic background of lineage IV (Figure 5 and Supplementary Figure 7), a high-risk clone characterized by the presence of *bla*_NDM-1_ within plasmid p14ARS_VSM0843-1 displayed local clonal expansion in one sentinel site, while another high-risk clone characterized by the presence of *bla*_NDM-1_ within plasmid p13ARS_GMH0099 had also disseminated geographically across five different sites (Figure 5). Clonal expansion followed by geographical dissemination is usually attributed to the acquisition of an AMR determinant ^36^. The extent of the geographical dissemination of the different high-risk clones could be attributed to the different plasmid backbones (vehicle), but differences in the genetic background (strain) could also be at play. The presence of a robust AMR surveillance network such as the ARSP is key for the detection of geographical dissemination of high-risk clones at a regional/national level, and for the dissemination of the information to national stakeholders. In this respect, the ARSRL is currently evaluating formats for the inclusion of genomic data into their annual surveillance reports. The reference laboratory directors can also alert hospitals across the country to establish concerted infection control measures, as well as relay this information to the national Department of Health for the formulation of evidence-based guidelines.

At the global scale, we identified several high-risk clones of *E. coli* ST410 circulating the in the Philippines, with evidence of frequent international transmission, in agreement with a previous report ^18^. Previous reports of carbapenem-resistant *E. coli* ST410 from the Western Pacific and South East Asia regions described the presence of *bla*_OXA-181_ in China ^17, 37^, *bla*_NDM-1_ in Singapore ^38^, *bla*_NDM-5_ in South Korea ^39^, or the combination of *bla*_NDM-5_ and *bla*_OXA-181_ in Myanmar ^40^ and South Korea ^39^. The repertoire of carbapenemases in the retrospective collection of ST410 from the Philippines seemed to be in tune with the genes/variants circulating locally, as most isolates carried the prevalent variant *bla*_NDM-1_. Of note was the high-risk clone carrying both *bla*_NDM-1_ and *bla*_OXA-181_, a combination hitherto unreported in *E. coli* ST410, and acquired through the combination of a plasmid of local circulation and a conserved international plasmid, respectively. This suggests that *E. coli* ST410, a successful pandemic lineage, can not only easily disseminate, but it can also adjust the complement of carbapenemases by acquiring endemic plasmids along the way. Global AMR surveillance networks, such as the WHO GLASS ^10^, are paramount to detect the emergence and monitor the spread of resistance at the international level and inform the implementation of targeted prevention and control programmes. The ARSRL has been supplying laboratory surveillance data to GLASS since 2015, and presented their experience with WGS at a WHO technical consultation on the application of WGS for national, regional and global surveillance in 2020, thus becoming an international contributor in both fields.

Collecting information on AMR that can be rapidly transformed into action requires harmonized standards, especially at the national and international levels ^10^. For laboratory data, WHONET serves this purpose in the ARSP and in over 2,000 clinical, public health, veterinary, and food laboratories in over 120 countries worldwide ^26^, while GLASS serves this purpose at the global level. Genomic data are amenable to standardization ^29^, and national public health agencies ^41^ and international surveillance networks ^42^ are adopting diverse schema to identify and name genetic lineages. However, a global standard system for defining a cluster has not been implemented beyond the level of discrimination of MLST. Likewise, different databases of AMR mechanisms ^43–45^ may differ in content and nomenclature. Thus, the standardization of genomic data, as well as the provision of platforms for the uptake of genomic information are crucial moving forward, and for ongoing AMR surveillance.

Containment of AMR at a global level requires an international concerted effort. High-risk clones have the propensity to disseminate rapidly, and genomics can improve the detection of their emergence and spread. This highlights the increasing need to build equitable partnerships to facilitate ownership transfer of genomic epidemiology capacity (operational, analytical and interpretational) to enhance national AMR surveillance programmes within low-resource settings. The binational partnership of the See and Sequence project led to a large retrospective survey of bacterial pathogens that has provided the first in-depth genomic view of the AMR landscape in the Philippines and established contextual data for ongoing local prospective sequencing. In parallel, through training and transfer of expertise in laboratory procedures and bioinformatics, the open exchange of data, and collective interpretation of results, we have expanded the existing capacity of a national reference laboratory with WGS focused on action for public health. Whole genome sequencing commenced at the ARSRL in 2018 with the Illumina MiSeq equipment available locally. A new dedicated bioinformatics server was installed at ARSRL for sequence data storage and analysis. Collective data interpretation was aided by data sharing via interactive web tools such as Microreact (www.microreact.org ^27^) and Pathogenwatch (www.pathogen.watch) to leverage the expertise from both CGPS and ARSRL. The results of the retrospective survey were presented to representatives of the sentinel sites during the ARSP annual meetings and at international conferences, at first by members of the CGPS, followed by presentations by members of the ARSRL in subsequent years ^46, 47^. The ARSRL staff conducted its first outbreak investigation using WGS in July 2019, with sequencing, bioinformatics, and reporting to the DOH conducted locally, with a turnaround time of eight days from sample receipt to report preparation. While progress has been made, significant challenges remain to bring genomics into routine surveillance in low-resource settings. These include, and are not limited to, challenges in the supply chain and procurement of WGS reagents and equipment, vastly differing costs between high- and low-income settings, shortage of skilled local bioinformaticians due to both limited access to training and difficulties in staff retention, and the lack of platforms to feedback the actionable genomic data to sentinel sites. In addition, it was expected that the timeline from implementation of a disruptive technology such as WGS to full adoption by all the relevant stakeholders was to exceed the period of one academically-funded project. The continued collaboration with the ARSRL (via an NIHR-funded project) is building on the foundations laid by See and Sequence to tackle some of the challenges mentioned above, and to improve on the translation of genomic data into public-health action and policy. We generated a blueprint for the sustainable implementation of WGS and genomic epidemiology within national surveillance networks in LMICs that can be adapted and utilized within other locations to tackle the global challenge of AMR.

## Methods

### ARSP workflow and data

The ARSP implements antimicrobial resistance surveillance on aerobic bacteria from clinical specimens. The program collects culture and antimicrobial susceptibility data from its 24 sentinel sites and 2 gonorrhoeae surveillance sites. The sentinel sites participate in an external quality assessment scheme (EQAS) conducted by the reference laboratory to ensure quality of laboratory results. Case-finding is based on priority specimens sent routinely to the hospitals’ laboratories for clinical purposes, and is thus based on the diagnostic practices of the clinicians, as well as the resources available to the patient to cover the culture and susceptibility testing. Sentinel sites submit monthly results of all culture and susceptibility tests to the Antibiotic Resistance Surveillance Reference Laboratory (ARSRL) via WHONET, a free Windows-based database software for the management and analysis of microbiology laboratory data with a special focus on the analysis of antimicrobial susceptibility test results ^8^. Multi-drug resistant phenotypes as classified according to recommended standard definitions ^48^. Select isolates are referred to the ARSRL for confirmatory testing if they fall in one of the three following categories: i) organisms of public-health importance regardless of their resistance phenotype, such as *Salmonella enterica* ser. Typhi, *Streptococcus pneumoniae* and *Vibrio cholerae*; ii) organisms with emerging resistance, such as carbapenem-resistance; iii) organisms with rare susceptibility patterns, such as vancomycin-resistant *Staphylococcus aureus* or ceftriaxone-resistant *Neisseria gonorrhoeae*. At the ARSRL, all referred isolates are re-identified using standardized culture, and susceptibility tested by disc diffusion or minimum inhibitory concentration (MIC) determined using the most current guidance for antibiotic test panels and breakpoints recommended by the Clinical and Laboratory Standards Institute (CLSI). Serotyping for *S. pneumoniae, H. influenzae*, Salmonellae and *V. cholerae* is conducted to further characterize the isolates.

### ARSP data and bacterial isolates included in this study

Phenotypic data collected since 1988 from antimicrobial susceptibility tests (Supplementary Methods, Appendix) was summarized with WHONET to compute yearly resistance rates for key pathogen-antibiotic combinations. Only the first bacterial isolate per patient per species per year was included in the calculations.

Resistance profiles derived from the antibiograms were summarized per organism with WHONET ^8^, and their relative abundance visualized with Tableau Desktop. Data collected between 2009 and 2017 were included in the analysis as the number of sentinel sites collecting data remained relatively stable during this period (22-24 sites). Isolates with missing data for any antibiotic on the panels listed on Table 1 were excluded.

The bacterial pathogens included in the See and Sequence study were those responsible for the majority of antimicrobial resistant infections in the Philippines. These included carbapenemase-producing *Pseudomonas aeruginosa* and *Acinetobacter baumanii*, carbapenemase-producing and ESBL-suspect *Escherichia coli* and *Klebsiella pneumoniae*, methicillin-resistant *Staphylococcus aureus*, *Neisseria gonorrhoeae*, non-typhoidal *Salmonella*, and *Salmonella enterica* ser. Typhi. Linked epidemiological data included location and date of specimen collection, type of specimen, type of patient (in or outpatient), and sex and age of the patient. We utilized a proxy definition for “infection origin” whereby patient first isolates collected in the community or on either of the first two days of hospitalization were categorized as community infection isolates, while isolates collected on hospital day three or later were categorized as hospital-acquired infection isolates ^10^.

In this article we focus on carbapenemase-producing organisms. All carbapenemase-producing *K. pneumoniae* and *E. coli* isolates referred to, confirmed, and banked by the ARSRL in 2013-2014 were selected for the retrospective sequencing survey.

Approximately 100 isolates of carbapenemase-producing *P. aeruginosa* and *A. baumannii* each, were selected according to the following criteria: i) referred to ARSRL in 2013-2014; ii) complete resistance profile (i.e., no missing data); iii) overall prevalence of the resistance profile in the ARSP data (including both referred and non-referred isolates); iv) geographical representation of different sentinel sites. The number of isolates included from each sentinel site proportional to their relative abundance and estimated from (n/N)*100 (rounded up), where n is the total number of isolates from one site, and N is grand total of isolates ; v) when both invasive and non-invasive isolates representing a combination of resistance profile, sentinel site and year of collection were available, invasive isolates (i.e. from blood, or cerebrospinal, joint, pleural and pericardial fluids) were given priority. In addition, approximately 100 isolates of ESBL-producing *E. coli* and *K. pneumoniae* each were included as per the criteria above. Pan-susceptible isolates were only available for *P. aeruginosa* isolates and were also included in the retrospective sequencing survey.

Lyophilisates of the selected bacterial isolates were re-suspended in 0.5 mL of tryptic soy broth, plated onto MacConkey agar plates, and incubated at 35-37°C for 18-24 hours. The species identity of the revived bacterial isolates was confirmed by characterization of the colony morphology and conventional biochemical tests (e.g., oxidase, motility, and sugar fermentation using triple sugar iron agar test). Phenotypic re-testing was performed on the resuscitated isolates based on the phenotype stored in WHONET. Carbapenemase production in resuscitated isolates of Enterobacteriaceae was tested with the modified Hodge test (MHT), and confirmed using the imipenem MIC method with imipenem Etest strips (BioMerieux). *P. aeruginosa* and *A. baumanii* were screened for metallo-beta-lactamase production using the EDTA synergy test between imipenem disk (10ug) and EDTA (750ug/mL), which was confirmed by imipenem/imipenem inhibitor (IP/IPI) Etest strips. *E. coli* and *K. pneumoniae* isolates were screened for ESBL with the double-disk synergy test between aztreonam (30ug), and amoxicillin/clavulanic acid (20/10ug) and confirmed by using the gradient diffusion method with cefotaxime/cefotaxime with clavulanic acid (CT/CTL) or ceftazidime/ceftazidime with clavulanic acid (TZ/TZL) Etest strips.

### Whole Genome Sequencing, Assembly and Annotation

*A. baumanni*, *E. coli*, *K. pneumoniae*, and *P. aeruginosa* bacterial strains were shipped to the Wellcome Sanger Institute streaked onto Nutrient Agar butts contained in screw-cap cryovials, where DNA was extracted from a single colony of each isolate with the QIAamp 96 DNA QIAcube HT kit and a QIAcube HT (Qiagen; Hilden, Germany). DNA extracts were multiplexed and sequenced on the Illumina HiSeq platform (Illumina, CA, USA) with 100-bp paired-end reads.

The quality of the sequence data was assessed for median base quality >30, mapping coverage >80% of the length of a reference genome sequence of the same species (*A. baumannii* ATCC 17978, *E. coli* UPEC_ST131, *K. pneumoniae* subsp. *pneumoniae* Ecl8, and *P. aeruginosa* LESB58), and >80% of the sequence reads assigned to the corresponding species with Kraken v0.10.6-a2d113dc8f ^49^ and a reference a database of all viruses, archaea and bacteria genomes in RefSeq ^50^ and the mouse and human reference.

Annotated assemblies were produced using the pipeline described in ^51^. For each sample, sequence reads were used to create multiple assemblies using VelvetOptimiser v2.2.5 ^52^ and Velvet v1.2 ^53^. An assembly improvement step was applied to the assembly with the best N50 and contigs were scaffolded using SSPACE ^54^ and sequence gaps filled using GapFiller ^55^. Automated annotation was performed using PROKKA v1.5 ^56^ and a genus specific database from RefSeq ^50^. The quality of the assemblies was assessed for total size, number of contigs, N50, and GC content appropriate for each species, with all genomes passing quality control characterized by assemblies of <150 contigs, and N50 > 60,000.

### Phylogenetic analysis

Evolutionary relationships between isolates were inferred from single nucleotide polymorphisms (SNPs) by mapping the paired-end reads to reference genomes, using the Burrows Wheeler aligner (BWA) v0.7.12 ^57^ to produce a BAM file. PCR duplicate reads were identified using Picard v1.92 ^58^ and flagged as duplicates in the BAM file.

Variation detection was performed using samtools mpileup v0.1.19 ^59^ with parameters “-d 1000 -DSugBf” and bcftools v0.1.19 ^60^ to produce a BCF file of all variant sites. The option to call genotypes at variant sites was passed to the bcftools call. All bases were filtered to remove those with uncertainty in the base call. The bcftools variant quality score was required to be greater than 50 (quality < 50) and mapping quality greater than 30 (map_quality < 30). If not all reads gave the same base call, the allele frequency, as calculated by bcftools, was required to be either 0 for bases called the same as the reference, or 1 for bases called as a SNP (af1 < 0.95). The majority base call was required to be present in at least 75% of reads mapping at the base, (ratio < 0.75), and the minimum mapping depth required was 4 reads, at least two of which had to map to each strand (depth < 4, depth_strand < 2). Finally, strand_bias was required to be less than 0.001, map_bias less than 0.001 and tail_bias less than 0.001. If any of these filters were not met, the base was called as uncertain.

A pseudo-genome was constructed by substituting the base call at each site (variant and non-variant) in the BCF file into the reference genome and any site called as uncertain was substituted with an N. Insertions with respect to the reference genome were ignored and deletions with respect to the reference genome were filled with N’s in the pseudo-genome to keep it aligned and the same length as the reference genome used for read mapping. For sequence-type specific phylogenies, mobile genetic elements (MGEs) and recombination regions detected with Gubbins v1.4.10 ^61^ were masked from the alignment of pseudogenomes. Alignments of only variant positions were inferred with snp-sites v2.4.1 ^62^, and were also used to compute pairwise SNP differences between isolates with an in-house script.

Maximum-likelihood phylogenetic trees were generated with RAxML v8.2.8 ^63^ based on the generalised time reversible (GTR) model with GAMMA method of correction for among-site rate variation and 100 bootstrap replications. Trees were midpoint-rooted. The phylogenetic trees, genomic predictions of AMR, and genotyping results were visualized together with the ARSP epidemiological data using Microreact ^27^.

### In silico predictions of AMR determinants and plasmid replicons

Known antibiotic resistance determinants were identified using Antibiotic Resistance Identification By Assembly v.2.6.1 (ARIBA ^64^). For acquired genes an in-house curated database comprising 2316 known resistance gene variants was used (https://figshare.com/s/94437a301288969109c2) ^65^. Full-length assembled matches with identity >90% and coverage >5x were considered as evidence of the presence of the gene in the query genome. Further manual inspection of the results excluded 13 matches that passed the above filters but that were suspected low-level contamination. Mutations in *ompK35* and *ompK36* genes were detected with ARIBA and the Comprehensive Antimicrobial Resistance Database (CARD, ^44^) and the output was inspected for truncations, interruptions and frameshifts.

Plasmid replicons were identified with ARIBA and the PlasmidFinder database ^66^. Full-length assembled matches with 100% identity were considered as evidence of the presence of the replicon gene.

### In silico multi-locus sequence typing

Multi-locus sequence types (STs) were determined from assemblies with MLSTcheck v1.007001 ^67^ or from sequence reads with ARIBA ^64^, using species-specific databases hosted at PubMLST ^68^ for *E. coli ^69^*, *A. baumannii ^70^*, *P. aeruginosa ^71^*, or at BIGSdb ^68^ for *K. pneumoniae* ^72^.

MLST calls from *P. aeruginosa* genome sequences were manually curated as we noted the presence of two genes exhibiting 100% sequence identity to different *acsA* alleles in the pubmlst database (accessed April 2019), which confounded the *in silico* predictions. This was not privy to the genomes is this study, since the genome sequence of reference strain NCGM2_S1 (accession AP012280), known to belong to ST235, showed two genes with locus tags NCGM2_1665 (nt position 1808730-1810685) and NCGM2_0806 (nt positions c879796-881733) that are annotated as *acsA* and *acsB*, respectively, and are 100% identical to alleles acsA_38 and acsA_225, respectively (https://pubmlst.org/data/alleles/paeruginosa/acsA.tfa). Furthermore, the same was only observed for the *acsA* allele both from assemblies (MLST_check) and raw reads (ARIBA-MLST), and therefore unlikely the result of contamination with other strains. Allele acsA_38 defines ST235, but allele acsA_225 defines a novel ST. Thus, when two different *acsA* alleles were detected in a *P. aeruginosa* genome we selected the one associated with a known ST.

### Characterization of carbapenemase plasmids

Select *K. pneumoniae* (14ARS_CVM0040, 14ARS_MMH0055, 13ARS_GMH0099, 14ARS_VSM0843, 13ARS-VSM0593, and 13ARS_MMH0112) and *E. coli* (14ARS_NMC0074) isolates were also sequenced with PacBio RS II or Sequel (Pacific Biosciences, CA, USA) or Oxford Nanopore Gridion (Oxford Nanopore, Oxford, UK) platforms. Hybrid assemblies from short and long-reads were obtained with Unicycler v0.4.0 ^73^ with default parameters, except for 13ARS_GMH0099 for which the bold assembly mode was used. The sequence of the p13ARS_GMH0099 plasmid was manually curated with Bandage v0.8.1 ^74^ circularized with Circlator v1.5.3 ^75^, and further polished with Quiver (Pacific Biosciences, CA, USA).

The presence of AMR genes and plasmid replicons in the plasmid sequences was detected with Resfinder ^45^ and PlasmidFinder ^66^, respectively. Matches with identity larger than 98% were reported. Detailed annotations of plasmids were obtained with MARA ^76^. Visual comparisons between plasmid sequences were created with BRIG v1.0 ^77^ with default BLAST parameters.

The distribution of plasmids across draft genomes was inferred by mapping short reads to the plasmid sequences with smalt v0.7.4 with identity threshold y=96% and random alignment of reads with multiple mapping positions of equal score (-r 1). The BAM file was sorted and indexed with samtools v0.1.19-44428cd ^59^, which was also used to and generate a pileup (mpileup -DSugBf) of the alignment records in BCF format. A pseudo-plasmid sequence in fastq format was constructed with the bcftools view and vcfutils.pl vcf2fq programs, and the sequence length coverage (%) of the reference plasmid was computed.

## Acknowledgements

This work was funded by the Newton Fund, Medical Research Council (UK), Philippine Council for Health Research and Development, Centre for Genomic Pathogen Surveillance, and National Institute for Health Research (UK).

## Author Contributions

C.C. and D.M.A. conceived the study. The ARSP collected the bacterial isolates and epidemiological data and performed preliminary laboratory analyses. S.A., M.A.L.M., J.M.G., M.L.L., P.K.V.M., V.C., M.T.L., H.O.E., J.C.P., J.C., M.C.J., A.S.V., J.B.B., A.M.O., L.F.T.F., K.D.B., B.J., K.A., C.M.H., S.B.S., J.S., M.T.G.H, C.C., and D.M.A. performed the data analysis. S.A., M.A.L.M., M.T.G.H., C.C., and D.M.A. wrote the manuscript. All authors read and approved the manuscript.

Antimicrobial Resistance Surveillance Program: Jerry C. Abrogueña, Minda B. Aguenza, Sherine Alcantara, Evelyn M. Andamon, Myrna P. Angeles, Cecilia Belo, Julie B. Cabantog, Donna M. Calaoagan, Jobert A. Castillon, Joseph D. De Las Alas, Jerilyn L. Dulay, Mari Esguerra, Ann D. Fandida, Elizabeth Fangot, Hans D. Ferraris, Daisy Goco, Oscar P. Grageda, Bernadette B. Hapitana, Evelina Lagamayo, Modesty A. Leaño, Nena S. Lingayon, Myra Olicia, Rhodora B. Ongtangco, Jane G. Pagaddu, Aireen P. Parayno, Ingrid D. Peralta, Grace L. Pong, Cleocita P. Portula, Teresita N. Rebuldad, Maricel Ribo, Chanda G. Romero, Annette L. Salillas, Emerita T. Sinon, Karlo Tayzon, Edith S. Tria, Kristine Anne Vasquez, Januario D. Veloso, Ma. Merlina A. Vistal, Marilyn T. Zarraga

