## Supplementary Materials for "See and Sequence: Integrating Whole-Genome Sequencing Within the National Antimicrobial Resistance Surveillance Program in the Philippines"

### Supplementary Results and Figures

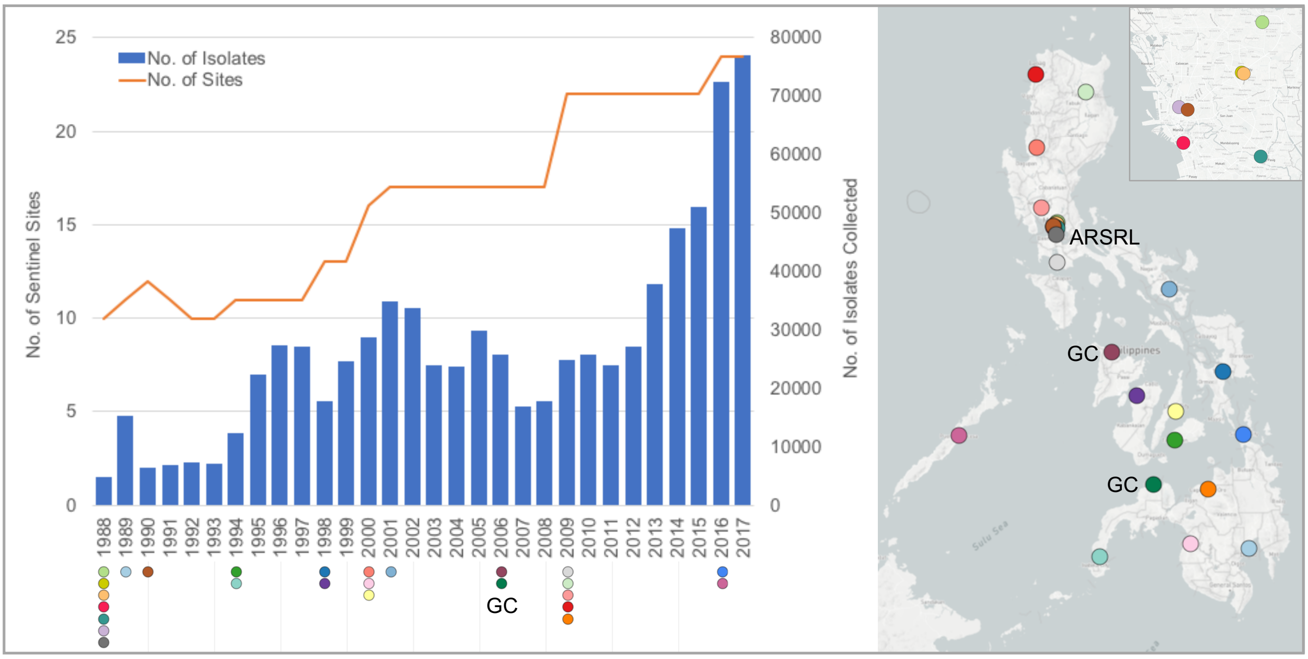


**Supplementary Figure S1. Three decades of antimicrobial resistance surveillance in the Philippines.** Total number of bacterial isolates collected by the sentinel sites per year (blue bars), number of sentinel sites participating in the program (orange line), their location (map) and year they joined ARSP (dots under the x-axis). The map inset shows the sentinel sites in the Metro Manila area. The number of sentinel sites does not include the 2 gonorrhea surveillance (GC) sites. Only the sites that have remained in the program until the time of publication are shown by coloured dots on the map and under the x-axis. ARSRL: reference lab.

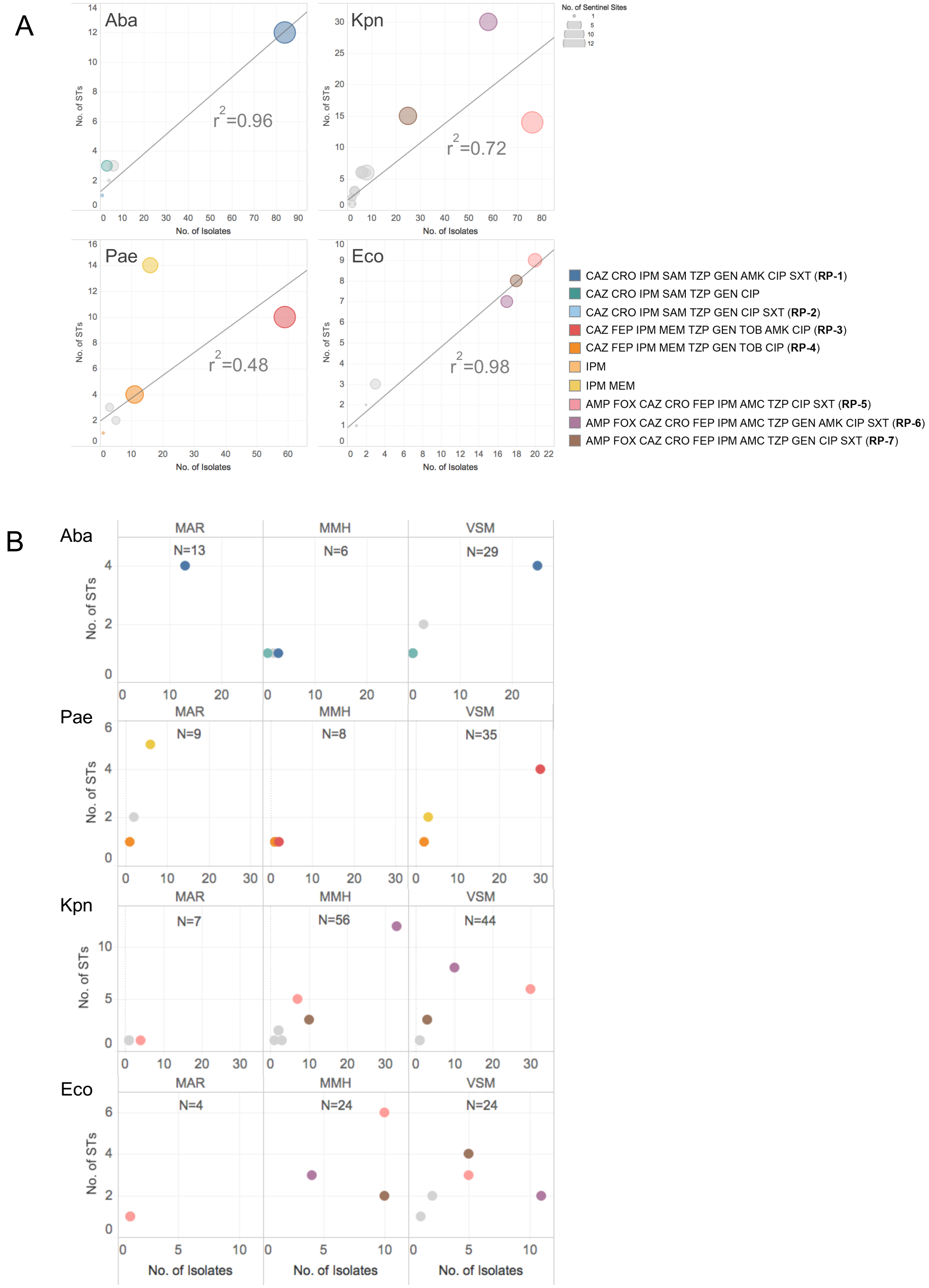


**Supplementary Figure S2. Relationship between carbapenem resistance profiles and distinct genetic lineages defined by their sequence type (ST).** Scatterplots portraying the number of STs as a function of the number of isolates sequenced retrospectively for the carbapenem resistance profiles. **A)** National level data (all sentinel sites combined). The size of the dot is proportional to the number of sentinel sites each profile was found on. **B)** Local-level data for the three sites contributing the largest number of isolates of the four organisms combined (MAR, MMH and VSM).

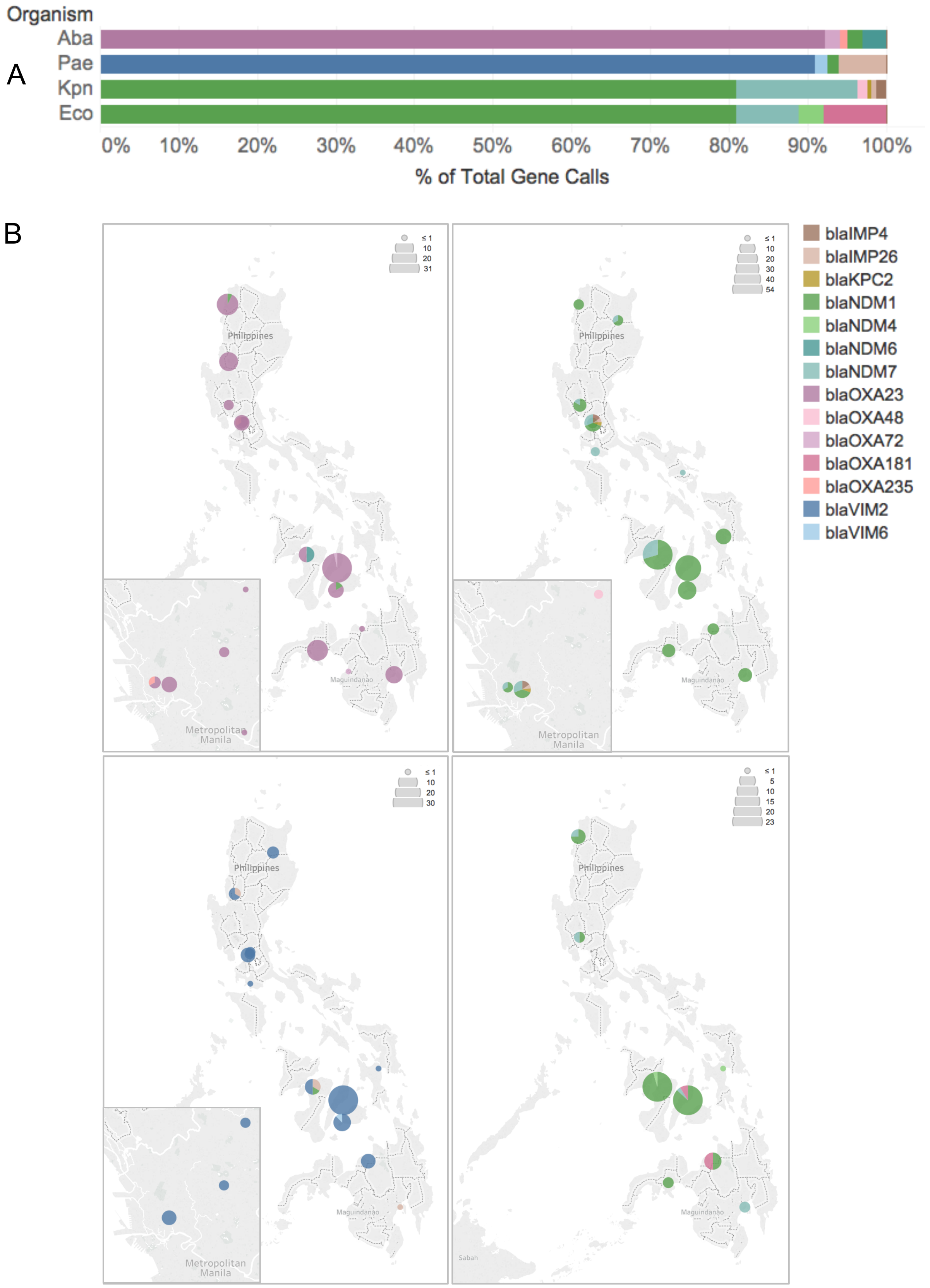


**Supplementary Figure S3. A)** Relative abundance of individual carbapenemase gene calls in *A. baumannii, P. aeruginosa, K. pneumoniae*, and *E. coli* genomes. **B)** Geographic distribution and relative abundance of carbapenemase genes across the sentinel sites. The size of the piecharts is proportional to the total number of individual gene calls for each organism. Map insets show the detail of the national capital region.

*Scale of surveillance – Local: WGS reveals a plasmid driven hospital outbreak of carbapenem-resistant K. pneumoniae*

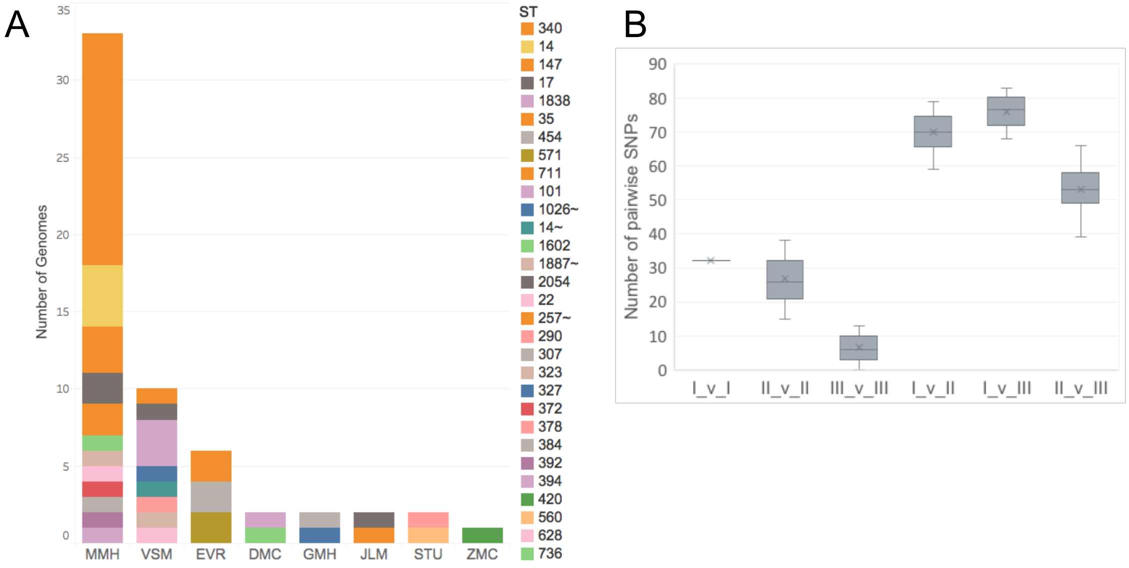


**Supplementary Figure S4. A)** Breakdown of *K. pneumoniae* genomes with resistance phenotype "AMP FOX CAZ CRO FEP IPM AMC TZP GEN AMK CIP SXT" by sentinel site and ST. **B)** Box and whisker plot of pairwise SNP differences within and between the three clades defined in Figure 4A. The median is indicated by the horizontal line, while the mean is indicated by an x. The whiskers indicate variability outside the upper and lower quartiles (box bounds). The pairwise SNP differences between genomes from clade III (N=105 pairwise comparisons) are significantly lower than those between genomes from clade II (N=15 pairwise comparisons, two-tailed Mann-Whitney U test, z-score -6.265, p-value = 3.70767e-10).

Plasmid p13ARS_MMH0112-3 exhibited high sequence similarity to three plasmids found in international isolates of other Enterobacteriaceae, as well as to plasmid p14ARS_CVM0040-2 from a *K. pneumoniae* ST147 isolate from this study (Supplementary Figure S5B). The main difference was the lack of a transposon carrying KpsC and KpsS, both encoding capsule polysaccharide biosynthesis proteins, in p13ARS_MMH0112-3. This suggests p13ARS_MMH0112-3 may have derived from an epidemic plasmid of global circulation that also circulates in the Philippines.

A second large plasmid (p13ARS_MMH0112-2) carried the ESBL gene *bla*$_{\text{CTX-M-15}}$, as well as other AMR genes and genomic islands for ferric dicitrate transport and resistance to heavy metals (Table 2 and Supplementary Figure S5A). Resistance to heavy-metals can aid survival in the hospital environment (1), while iron acquisition is crucial for survival within the host (2). By mapping short Illumina reads of all 33 possible-XDR genomes from MMH to the p13ARS_MMH0112-2 sequence, we found the entire plasmid sequence (i.e., ≥95% of the length) represented only in the 15 ST340 genomes, suggesting that this plasmid contributed to the persistence of this clone in the hospital environment, and in particular in the NICU (Supplementary Figure S5C). This is of significance in view of the non-overlapping hospitalization periods for most of the NICU patients.

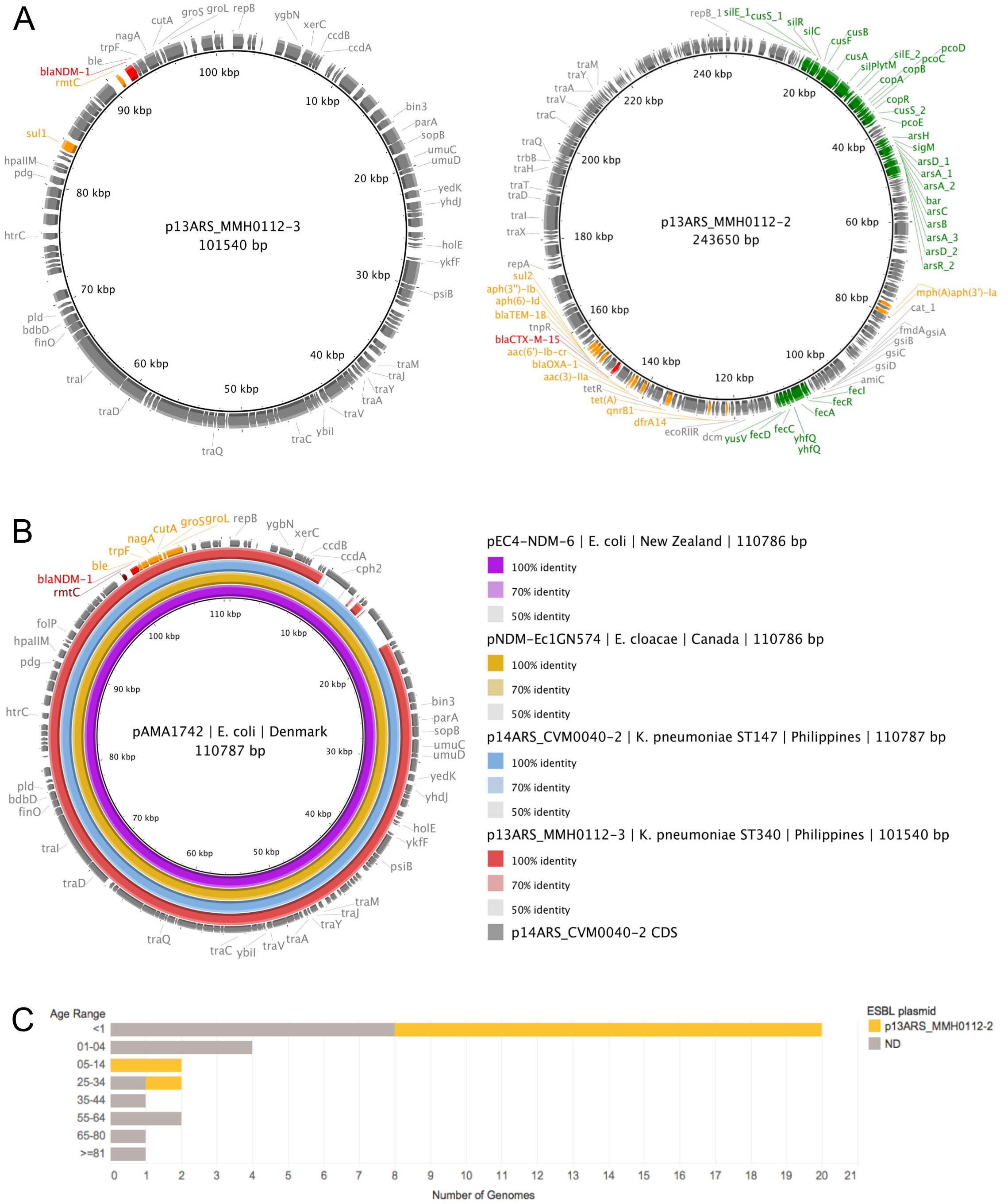

**Supplementary Figure S5. Plasmids from *K. pneumoniae* ST340 strain 13ARS_MMH0112. A)** Gene map highlighting the resistance genes (*bla*$_{NDM-1}$ and *bla*$_{CTX-M-15}$ in red and the rest in orange), and genomic islands of heavy metal resistance and iron acquisition (green). **B)** Comparison of p13ARS_MMH0112-3 to three international plasmids and to p14ARSCVM0040-2 from this study using BRIG. Plasmid pAMA1742 was used as a reference (innermost black circle). **C)** Distribution of plasmid p13ARS_MMH0112-2 carrying *bla*$_{CTX-M-15}$ (yellow) across the 33 retrospective isolates from hospital MMH with resistance profile "AMP FOX CAZ CRO FEP IPM AMC TZP GEN AMK CIP SXT" broken down by patient age group. Short reads of the 33 retrospective isolates were mapped to the p13ARS_MMH0112-2 sequence and a plasmid match was accounted when the reads covered at least 95% of the sequence length with at least 5x depth of coverage. ND: p13ARS_MMH0112-2 not detected.

*Scale of surveillance - National: WGS reveals the interplay of carbapenem resistance genes and plasmids in the regional circulation of a successful K. pneumoniae lineage*

The tree of 160 ST147 global genomes shows a wide diversity of carbapenemase genes and variants within this sequence type (Supplementary Figures S6A). The Philippine genomes from clades I-III are interspersed with international genomes, suggesting that they represent lineages of international circulation. For example, the genomes in clade III-B cluster with seven genomes from Singapore (N=4), Poland (N=2), Germany (N=1) and Denmark (N=1), from which they differ by an average of 63 (+/-10) pairwise SNPs. On the other hand, the genomes in clade IV are more distantly related to the global genomes, differing from the closest genomes from Singapore (N=1) and Turkey (N=5) by an average of 114 (+/- 7) SNPs. This observation together with the lack of matches to plasmids p13ARS_GMH0099 and p14ARS_VSM0843-1 in the nucleotide databases, and the broader geographic distribution of clade IV in the Philippines, suggests that it may be unique to this country, though a larger representation of ST147 genomes from Asia is needed to ascertain this.

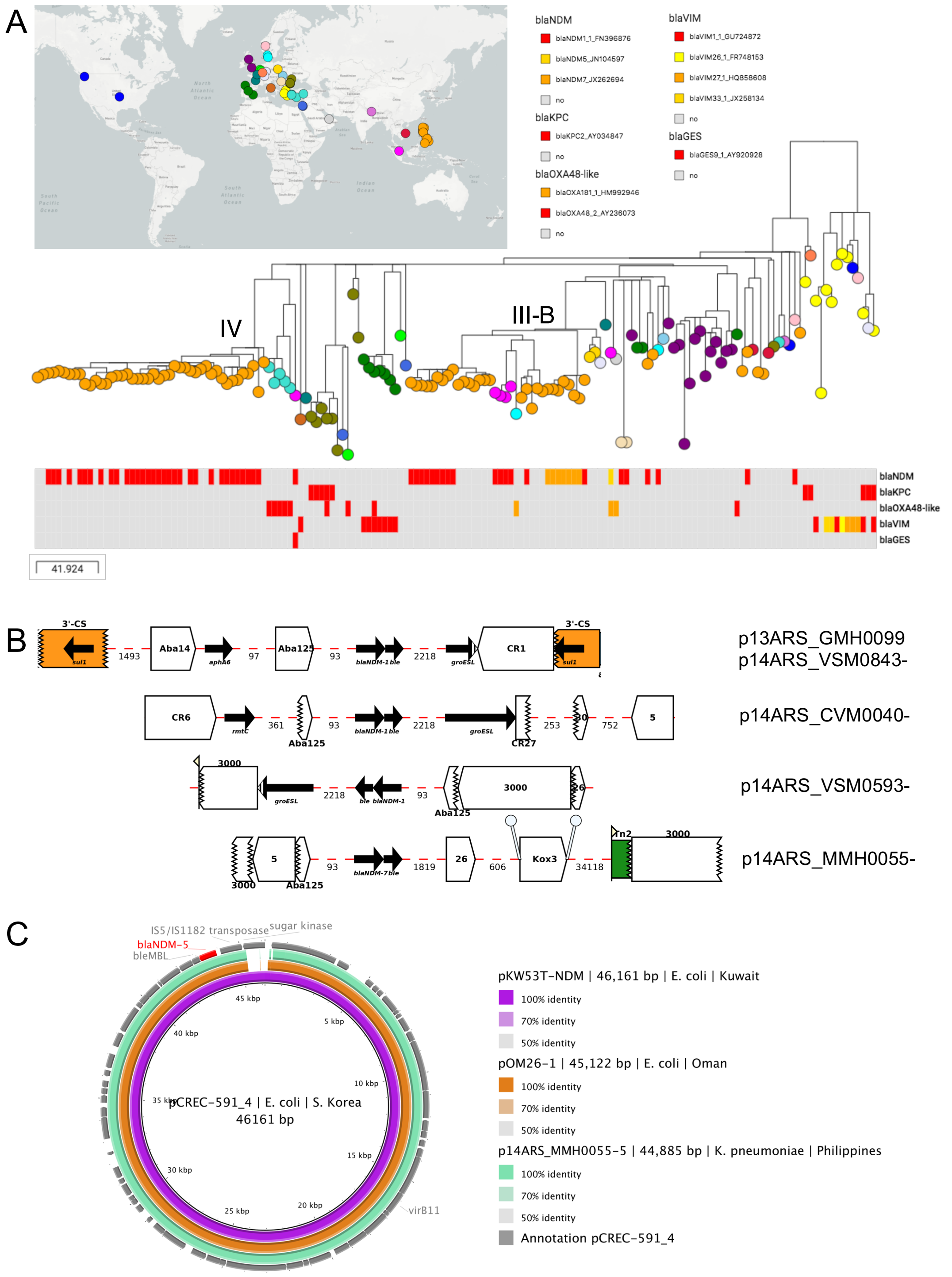


**Supplementary Figure S6. A)** Phylogenetic analysis of 80 ST147 genomes from the Philippines in global context. This interactive view is available at https://microreact.org/project/ARSP_KPN_ST147_GLOBAL/f21f2b91. The maximum-likelihood tree of 160 genomes was inferred from 3367 SNP sites identified from mapping the genomes to reference MS6671 (LN824133.1), after regions of MGEs and recombination were masked. The data are available at https://microreact.org/project/ARSP_KPN_ST147_GLOBAL. The distribution of carbapenemase genes and variants is shown as blocks under the tree. **B)** Context of NDM genes showing either intact (solid lines) or truncated (jagged lines) IS*Aba125* upstream elements, as annotated by MARA. **C)** Comparison of p14ARS_MMH0055-5 to three international plasmids (CP024825.1, KX214669.1, and KP776609.1) using BRIG. Plasmid pCREC-591_4 was used as a reference (innermost black circle).

We did not identify an NDM gene (or any other carbapenemase) in 11 genomes from carbapenem non-susceptible ST147 isolates (Figure 5). Resistance could not be explained by the combinatorial mechanism of ESBL gene (blaCTX-M-15) and OmpK poring disruption, as both the *ompk35* and *ompk36* genes were intact in these genomes (Supplementary Figure S7). Notably, the 11 genomes were found within the subclade characterized by the presence of the large mosaic plasmid p13ARS_GMH0099. The distribution of AMR genes showed that the genomes harboured most of the Tn2 transposon that carries *bla*$_{TEM-1}$ interrupted by the IS*Ecp1*-*bla*$_{CTX-M-15}$ element but had lost most of the remaining AMR genes (Supplementary Figure S7). IS*Ecp*1 usually mobilizes the ESBL gene, and has been reported to trigger transposition events to the chromosome (3). However, we could not confirm a chromosomal location in the draft assemblies, as the contigs containing *bla*$_{CTX-M-15}$ and *bla*$_{TEM-1}$ did not span beyond the flanking transposases. Alternatively, plasmid damage or partial plasmid loss may have occurred between resuscitation of the isolates and WGS.

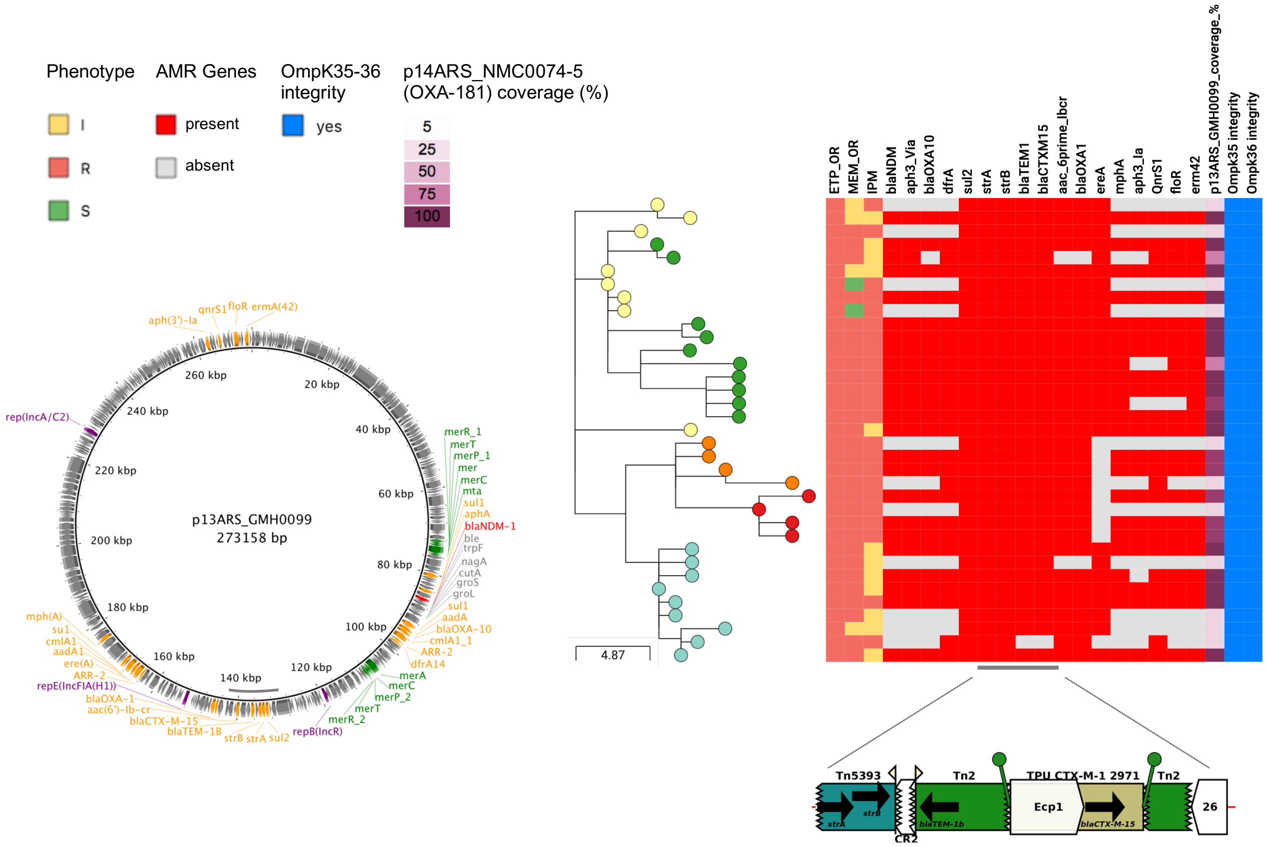


**Supplementary Figure S7. Potential NDM-1 plasmid loss within clade IV.** Distribution of AMR genes and of plasmid p13ARS_GMH0099 inferred from the short reads of genomes in clade IV. The AMR genes are displayed as per the order in the plasmid, and genes present in multiple copies were omitted. The detailed structure of Tn2-IS*Ecp*1 is shown below the gene blocks, and highlighted by a grey bar.

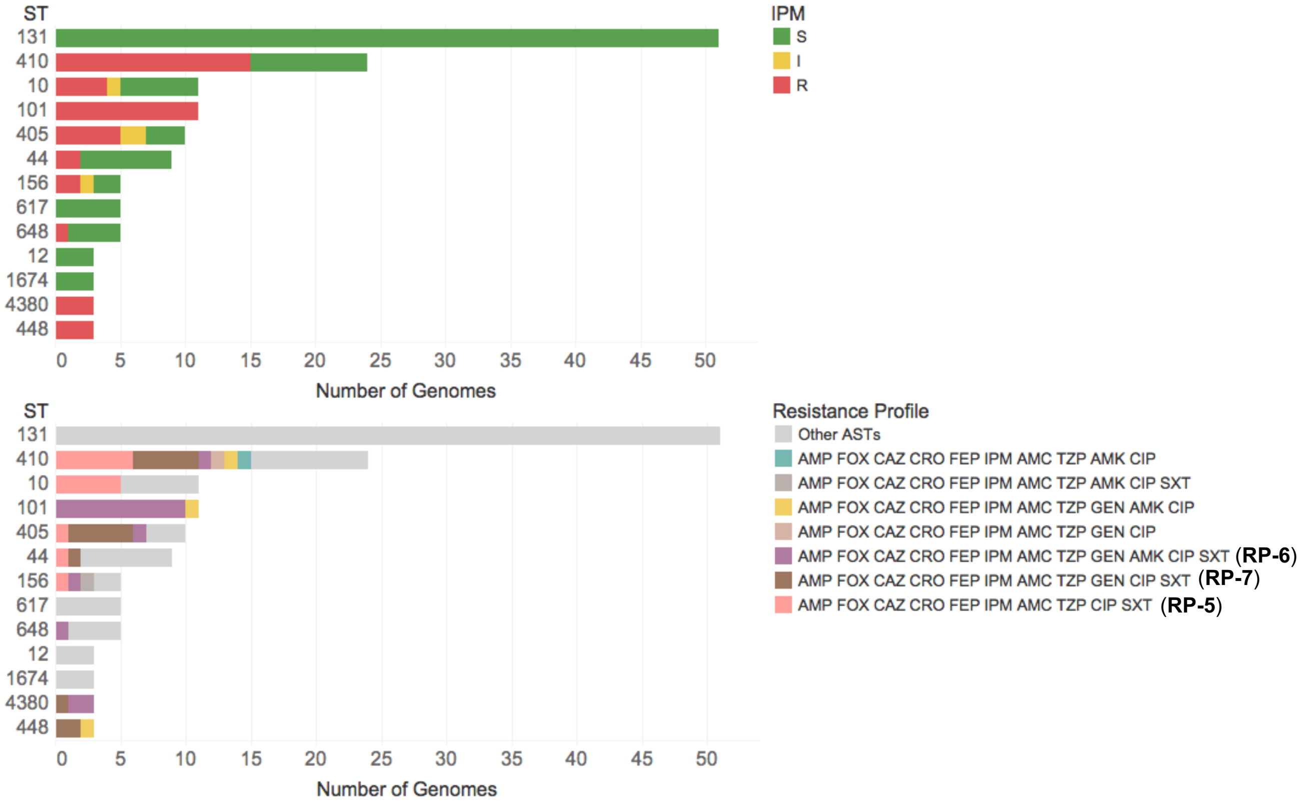


**Supplementary Figure S8.** Distribution of resistance to imipenem (top) and resistance profiles (bottom) across the *E. coli* STs. Only STs represented by at least three genomes are shown. S: susceptible; I: intermediate; R: resistant. Other ASTs: any other resistance profile with susceptibility to imipenem.

The phylogenetic analysis of *E. coli* ST410 genomes in global context showed that the three clones with possible-XDR isolates are included within a major lineage delineated by the presence of *bla*$_{CMY-2}$, and characterized by the presence of a larger number of resistance genes per genome (mean number 16·31 ± 4·85, N=44, vs 7·45 ± 2·54, N=36, Mann-Whitney U-test z score 6.824, $p$= 8.86846x10$^{-12}$), fewer genetic differences between genomes (mean number of pairwise SNP differences 41·55 ± 36·72, N=665 pairwise comparisons, vs 157·98 ± 47·68, N=989 pairwise comparisons, two-tailed Mann-Whitney U-test z score= -31.23, $p$< 0·0001), the presence of clinical and environmental isolates, and global distribution (Figure 6B), consistent with the characteristics of high-risk clones.

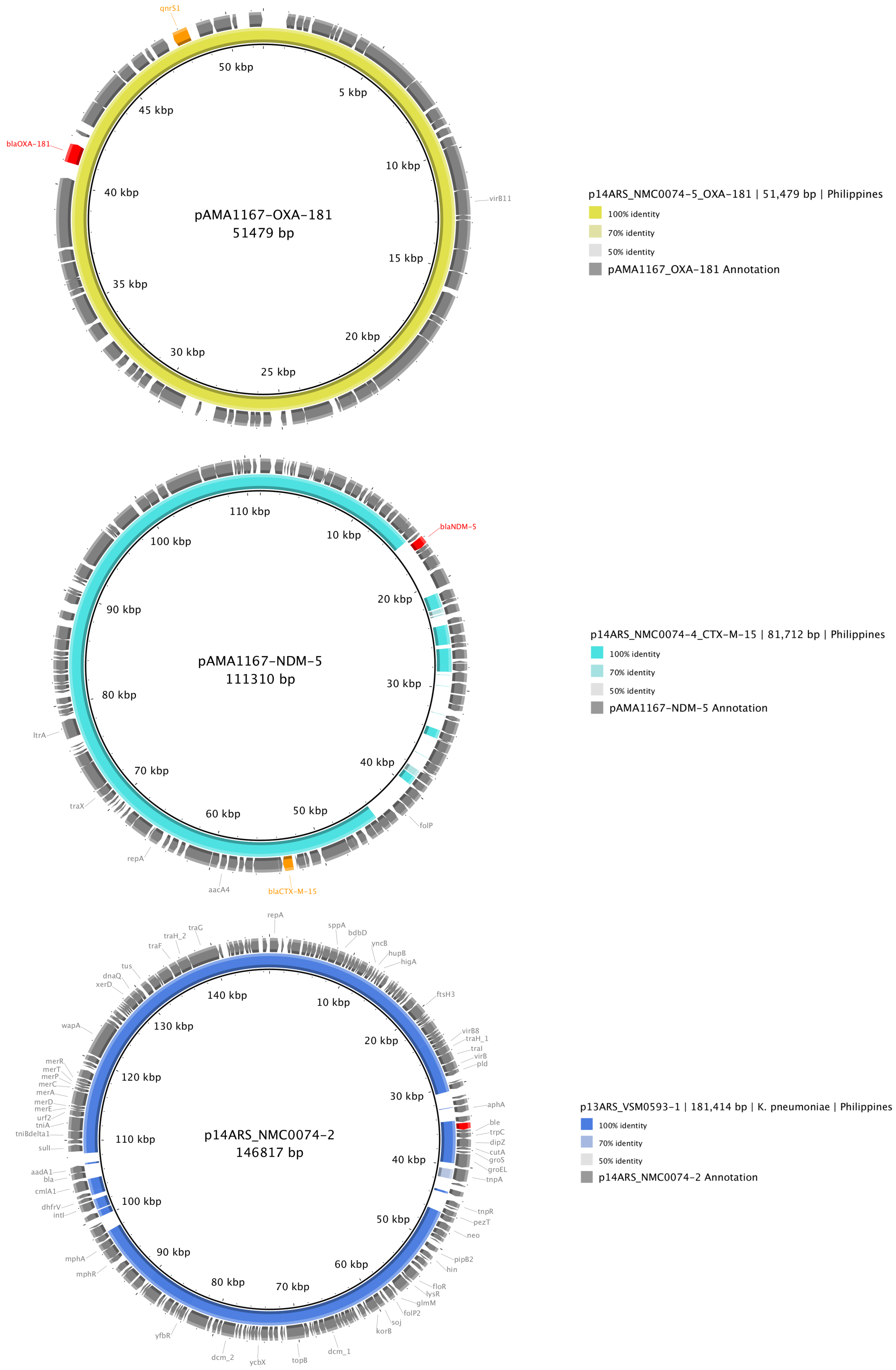


**Supplementary Figure S9. Plasmids from *E. coli* ST410 strain 14ARS_NMC0074 carrying two carbapenemase genes.** Top: Comparison of plasmid p14ARS_NMC0074-5 carrying $bla_{\text{OXA-181}}$ to plasmid pAMA1167-OXA-181 (CP024806.1). Middle: Comparison of plasmid p14ARS_NMC0074-4 carrying $bla_{\text{CTX-M-15}}$ to pAMA1167-NDM-5 (CP024805.1). Bottom: Comparison of plasmid p14ARS_NMC0074-2 carrying $bla_{\text{NDM-1}}$ to p13ARS_VSM0593-1 from a *K. pneumoniae* ST147 strain from this study.
